# Large-scale endoplasmic reticulum membrane solidification spatially organises proteins under thermal or metabolic stress

**DOI:** 10.64898/2026.04.09.717431

**Authors:** Paul M. Müller, Melissa R. Mikolaj, Elmehdi Belbaraka, Finn Hartstein, Sophia Altinoluk, Björn Perder, Philipp Trnka, Robert-William Welke, Heike Naumann, Nina Taudien, Severine Kunz, Michele Solimena, Ilya Levental, Kandice R. Levental, Andreas Müller, Helge Ewers, Kedar Narayan, Oliver Rocks

## Abstract

Organelle homeostasis is a key determinant of cellular fitness, yet how cells remodel their membranes in response to environmental change remains unclear. Here, we identify a temperature- and lipid saturation-dependent transformation of endoplasmic reticulum membranes into giant, rigid, multilamellar tubes in cells and *in vivo*. These ‘rods’ emerge from demixing of saturated lipids into solid-like domains – a previously unrecognised, large-scale endomembrane phase behaviour, fundamentally distinct from the transient liquid-ordered nanodomains of the plasma membrane. ER-tubulating reticulon-homology proteins are excluded from rods; their segregation drives progressive membrane flattening and ultimately multilayered wrapping. Surfactant-producing alveolar type-II lung cells, enriched in saturated lipids, form rods even at 37°C, demonstrating that native lipid metabolism can induce this transformation. This spatially organizing lipid-protein domain interplay may tune the ER tubule/sheet balance and provide a homeoviscous mechanism to preserve fluidity in the cholesterol-poor ER under thermal or metabolic stress.

## Introduction

Eukaryotic cells continuously remodel their membrane architecture and composition to organise biochemical processes and adapt to environmental changes. The endoplasmic reticulum (ER) is the most structurally and functionally versatile organelle, comprising the nuclear envelope and a dynamic network of tubules with further nanoscale morphologies^1–4^. ER sheets are major sites of protein biosynthesis, while tubules mediate lipid metabolism, Ca²⁺-signalling and inter-organelle communication, together forming a central synthesis, processing and distribution hub^5,6^. Cellular homeostasis depends on balanced ER structure and function, yet how the ER adapts to metabolic cues or stress remains incompletely understood^7^. Loss of ER integrity disrupts proteostasis and triggers the unfolded protein response implicated in numerous diseases^8^. Conversely, cancer cells exploit ER plasticity to sustain anabolic growth and survival under adverse conditions^9^.

At the molecular level, ER architecture is governed by its lipid composition, membrane-shaping proteins and cytoskeletal inputs^5,6^. Tubule-promoting reticulon-homology proteins insert paired hydrophobic hairpins into the cytosolic leaflet to induce curvature^10–14^, while CLIMP-63 stabilises sheets through luminal dimerisation^15,16^. By contrast, how the lipid matrix contributes to ER form and function is less well understood. Unlike the plasma membrane, which contains cholesterol-and sphingolipid-enriched liquid-ordered nanodomains, often termed ‘rafts’, that are associated with increased local bilayer stiffness and barrier properties, the ER is enriched in unsaturated phospholipids and contains only minimal cholesterol and sphingolipids^17–19^. This composition yields a loosely packed, liquid-disordered bilayer that facilitates bending and remodelling – even though ordered subdomains can form in yeast, and in mammals upon extreme perturbation^20–25^. A persistent challenge for such a fluid membrane is coping with temperature fluctuations and with metabolic changes that alter the ratio of saturated to unsaturated fatty acids (SFA/UFA). Cooling or SFA accumulation promotes tight lipid packing and potentially the formation of solid-like/gel lipid phases. Cells counter these effects through cholesterol^26^, and by adjusting the SFA/UFA synthesis, a process termed homeoviscous adaptation^27–29^. While the consequences of temperature and lipid composition on membrane biophysics are well established *in vitro*, their impact on organelle compartmentalisation and morphology in living cells is largely unexplored.

Here, we show that both hypothermia and increased lipid saturation induce a striking reorganisation of ER membranes into giant, rigid, multilamellar tubular structures (‘rods’) driven by large-scale demixing of solid-like lipid domains and segregation of ER-tubulating reticulon-homology proteins. These findings reveal fundamental principles of endomembrane phase behaviour, spatial organisation and homeoviscous balancing, and provide a tractable system to dissect phase coexistence and membrane plasticity under physical and metabolic stress.

## Results

### Temperature-dependent formation of rod-like structures from the endoplasmic reticulum

To investigate how temperature modulates organelle architecture, we expressed lipid-anchored fluorescent reporters in COS-7 cells. Unexpectedly, at room temperature (∼24°C), we observed remarkable long, straight membrane structures spanning the entire cell (Fig.1a). These ‘rods’ showed no preferential subcellular localisation and occasionally appeared interconnected. Rod staining was most pronounced with the hypervariable region (HVR) of the Rho GTPase CDC42, carrying a geranylgeranyl lipid anchor (Extended Data Fig.1a-c). Rods formed across all mammalian cell lines tested (COS-7, MDCK, HeLa, HEK293T, U-2 OS and C2C12 cells; Extended Data Fig.1d) and exhibited variable diameters, peaking at 0.6 µm in COS-7 cells (Fig.1c). Confocal z-stacks revealed a hollow tubular morphology (Fig.1b), explaining the paired-line appearance in single optical sections. BODIPY 493/503 staining confirmed their membranous nature: Pre- and post-cooling staining in untransfected cells demonstrated that rods form independent of reporter expression, transfection, or dye application (Extended Data Fig.1e).

**Fig.1.**
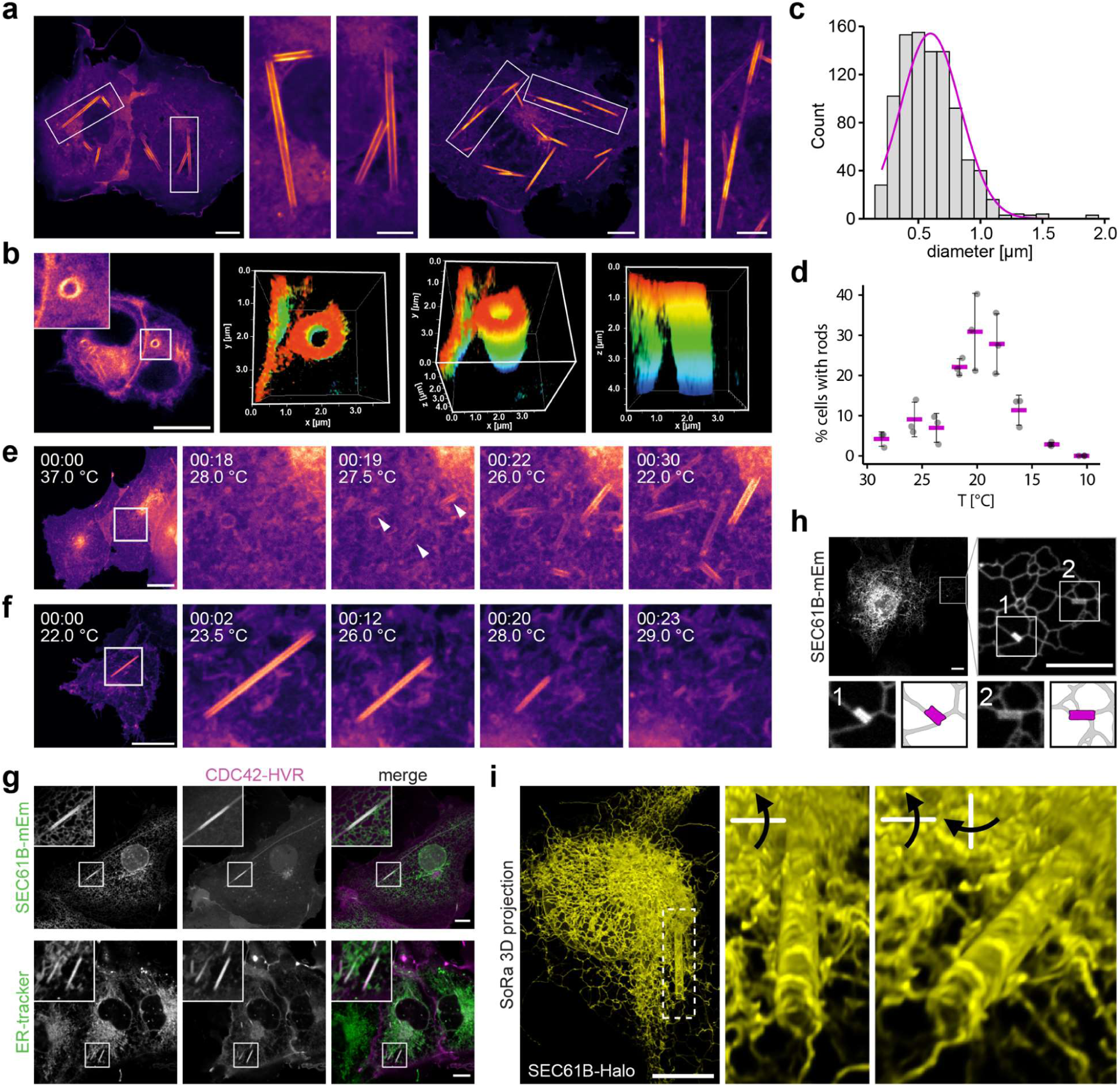
Temperature-dependent formation of rod-like structures from the ER. **a,** Confocal images of live COS-7 cells expressing the membrane marker mCitrine-CDC42-HVR after 40 min at 21°C. Insets show representative rod morphologies. Scale bars: 10 µm; insets: 5 µm. **b,** Left: Confocal image of a live HeLa cell expressing mCitrine-CDC42-HVR at 21°C. Right: 3D views of the boxed region (top view, 45°C tilted, side view; colour-coded in z). **c,** Histogram of rod diameters in COS-7 cells treated as in (**a**) (n=930). Magenta line: fitted Gaussian distribution. **d,** Rod formation in COS-7 cells expressing CDC42-HVR after 40 min at the indicated temperatures. Magenta crossbars: mean of three independent experiments (grey dots, 69-115 cells per experiment). Error bars: SD. **e,** Time-lapse confocal images of COS-7 cells expressing mCitrine-CDC42-HVR cooled from 37°C at 0.5°C/min using a Cherry Temp device. Arrowheads mark early rod structures. **f,** Time-lapse imaging of cells treated as in (**a**) during warming from 22°C at 0.25°C/min. **g,** Confocal live images of COS-7 cells expressing the ER marker SEC61B-mEm together with mCherry-CDC42-HVR (corresponds to Supplementary Video5), or mCitrine-CDC42-HVR co-stained with ER-tracker Red, after 40 min at 21°C. **h,** Confocal live images of COS-7 cell expressing SEC61B-mEm after 40 min at 21°C. Insets showing nascent rods connected to ER tubules and schematic segmentation (rods: magenta, ER: grey). **i,** 3D projection from super-resolution spinning disk microscopy with optical pixel reassignment (SoRa; ∼1.4x resolution enhancement), of a live COS-7 cell expressing JF646-labelled SEC61B-Halo after cooling to 21°C. The dashed region is magnified and rotated on the right, arrows indicate direction of rotation (corresponds to Supplementary Video7). Scale bars: 10 µm.

Live imaging under controlled temperature conditions showed that rods appeared consistently below 30°C, with maximal formation upon cooling to 21°C (Fig.1d). They emerged rapidly, elongating bidirectionally to reach steady-state length within minutes (Fig.1e and Supplementary Video1), and persisted for hours at reduced temperatures without detectable cytotoxicity. Despite frequently moving and rotating in cells, rods remained stiff and linear, indicating high mechanical stability. Rewarming triggered shrinkage from both ends and disassembly within minutes (Fig.1f, Extended Data Fig.1f and Supplementary Videos2 and 3). Notably, cooling–warming cycles repeatedly induced formation and dissolution at different cellular locations, demonstrating full reversibility (Extended Data Fig.1g and Supplementary Video4) with no detectable impact on cell viability.

Rod formation was independent of cytoskeletal scaffolding or active remodelling. Rods showed no colocalisation with actin or tubulin, and disruption of cytoskeletal dynamics with cytochalasin D, nocodazole, or the ROCK inhibitor Y-27632 had no effect (Extended Data Fig.2a). Energy depletion with sodium azide and 2-deoxy-D-glucose also did not impair rod formation (Extended Data Fig.2b), arguing against ATP- or GTP-dependent mechanisms. Rod formation therefore is consistent with a passive, spontaneous biophysical process.

Rods consistently colocalised with ER markers (Fig.1g; Extended Data Fig.2c), while mitochondrial, Golgi, and early/late endosomal markers remained unaltered and distinct from rods (Extended Data Fig.2d). Live imaging of SEC61B–GFP and mCherry–CDC42–HVR revealed rapid remodelling of local ER into rods, with persistent colocalisation throughout their lifetime (Supplementary Video5). Nascent rods were continuous with ER tubules, both end-to-end and laterally (Fig.1h and Supplementary Video6). 3D super-resolution microscopy further resolved their open-ended, hollow architecture and direct connection to the ER (Fig.1i and Supplementary Video7).

Under stress, ER membranes can adopt tightly stacked ordered smooth ER (OSER) structures^30,31^. However, OSER induction^32^ at lower temperature did not promote rod formation, and OSER and rods remained morphologically distinct. Furthermore, cells with extensive OSER formation showed fewer rods, indicating that rods originate from regular ER membranes (Extended Data Fig.2e,f).

Collectively, these results reveal that cooling mammalian cells induces a reversible, passive reorganisation of ER membranes into giant, rigid tubular membrane structures.

### Ultrastructural analysis reveals a continuous multilamellar architecture of rods

To resolve the ultrastructure of rods, we employed correlative light and electron microscopy (CLEM) using fluorescently tagged CDC42-HVR and SEC61B. Following confocal imaging of fixed COS-7 cells, samples were processed for EM and individual rods were identified by correlating fluorescence signals with morphological landmarks. This approach enabled reliable visualisation of rods in longitudinal and cross-sectional orientations, confirming a rigid tubular morphology (Fig.2a). Identical structures were observed after high-pressure freezing and cryo-substitution, excluding fixation artefacts (Extended Data Fig.3a,b). Transmission EM showed that rods are open-ended and can engulf other cellular components, including mitochondria, ER membranes and cytoskeletal filaments (Fig.2b and Extended Data Fig.3a,b). Notably, rods consisted of multiple membrane layers (Fig.2b). The number of layers varied along their length and typically decreased towards the ends, consistent with the stepwise reduction in fluorescence intensity observed by confocal microscopy (Fig.1a and Extended Data Fig.3c).

**Fig.2.**
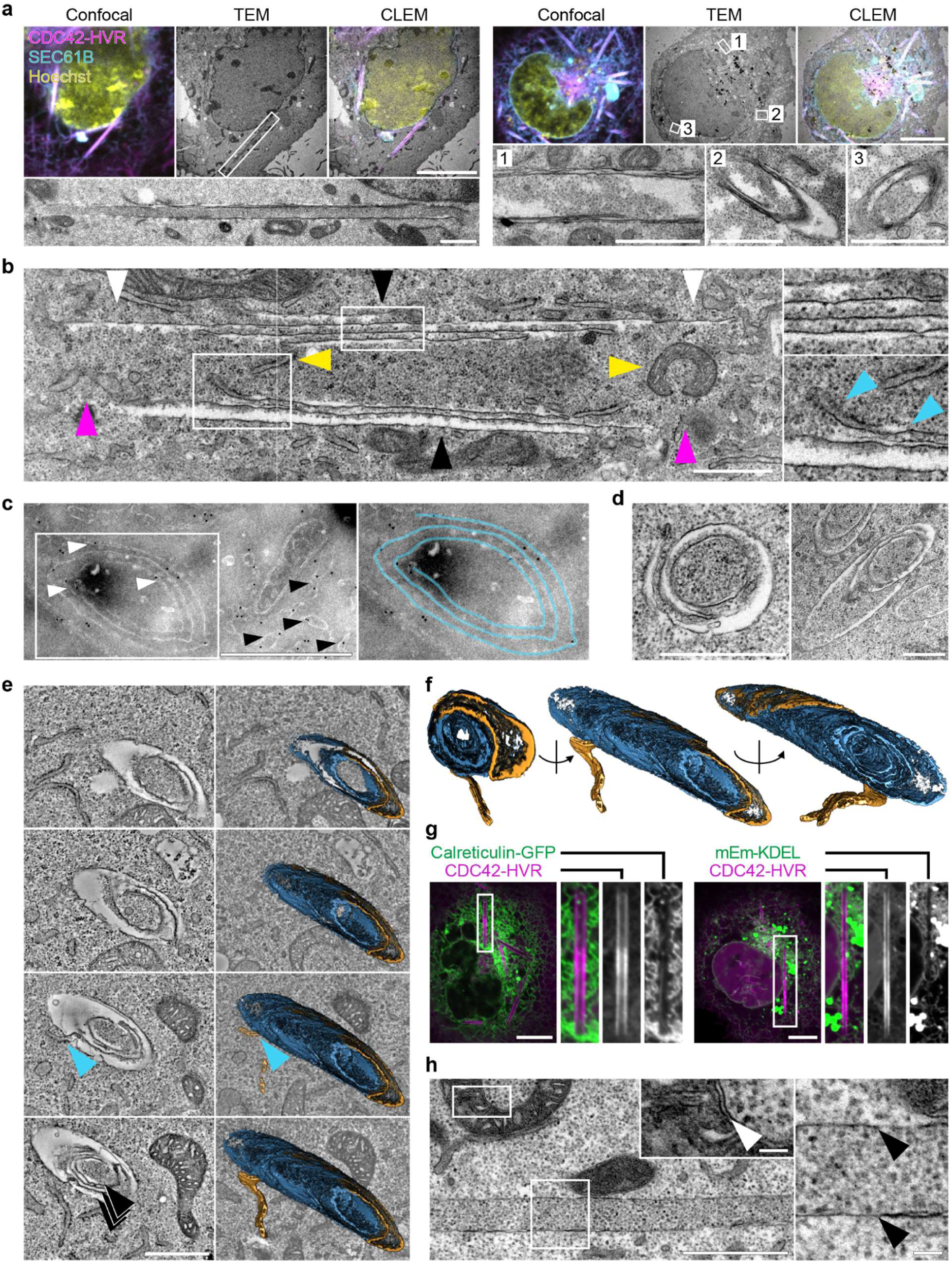
Ultrastructural analysis reveals multilamellar rod architecture and continuity with ER. **a,** CLEM images of COS-7 cells coexpressing mCherry-CDC42-HVR and SEC61B-mEm showing cross-sectional and longitudinal views of rods (TEM images of boxed regions are magnified below). Cells were equilibrated for 40 min at 21°C, stained with Hoechst, fixed, imaged by confocal microscopy and subsequently processed for TEM. Correlative mapping was performed using fluorescent landmarks. **b,** Longitudinal TEM section of rod revealing an open-ended architecture (magenta arrowheads), incorporation of small organelles (yellow arrowheads), membrane continuity between rods and the ER (bottom right, blue arrowheads), a multilamellar membrane organisation in the central region (top right and black arrowheads) and progressive tapering of layers towards the rod ends (black vs. white arrowheads). **c,** Immuno-EM of transfected COS-7 cells labelled with ø12nm-gold-conjugated antibodies against SEC61B-mEm. Immunogold particles localise to the ER (black arrowheads) and to multilamellar rod structures (white arrowheads), confirming the presence of ER proteins within rods. Right: Segmentation of the rod membrane showing spiral-like arrangement. **d,** TEM images of rod cross-sections confirming multilamellar spiral-like membrane topology. **e,** Left: Selected slices from serial-section electron tomograms of a rod in cross-section, revealing multilamellar organisation (black arrowheads) and membrane continuity between rods and ER (blue arrowheads). Right: 3D segmentation showing the rod membrane (blue) and the transition of connected ER membrane network (yellow). **f,** Through-the-hole view and side views of the segmented rod shown in (**e**). **g,** Representative spinning disk confocal micrographs of COS-7 cells expressing the luminal ER marker plasmids Calreticulin-GFP and mEmerald-KDEL, showing exclusion from rods relative to the rod marker mCherry-CDC42-HVR. **h,** TEM image of single layer rod lacking luminal space, in contrast to the intermembrane space of mitochondria. Both boxed regions are magnified at the same scale. Black arrowheads indicate rod membrane, white arrowhead indicates mitochondrial intermembrane space. Scale bars: 10 µm in (**a**) top row images and in (**g**); 0.1 µm in the magnified images in (**h**); 1 µm in all other images.

To resolve the three-dimensional membrane topology and its continuity with the surrounding ER, we performed serial-section electron tomography. While single sections already suggested a spiral-like arrangement of membrane layers (Fig.2c,d), tomographic reconstructions revealed a myelin-like architecture rather than separate concentric shells (Fig.2e,f and Supplementary Video8) and continuity between ER membranes and rods, both along the rod length and at rod ends(Fig.2e,f and Extended Data Fig.3d).

Consistent with this dense packing, confocal imaging showed a pronounced exclusion of the luminal ER proteins Calreticulin and the KDEL reporter from rods (Fig.2g; see also Fig.4a). This likely reflects physical exclusion of proteins with large luminal domains, as reported for highly constricted ER tubules (Hu et al. 2008). Accordingly, TEM of a rare single-layer rod revealed no discernible luminal space, in contrast to the clearly resolved intermembrane space of mitochondria in the same sections (Fig.2h). Together, these experiments demonstrate that rods are continuous, multilamellar, tightly packed membrane structures that are contiguous with the ER network.

### Rod formation is driven by lipid phase separation into solid-like domains

Because rod formation appeared spontaneous, we asked whether rods arise from lipid demixing in ER membranes at lower temperature. To compare membrane order between rods and the surrounding ER, we used the solvatochromic probe C-Laurdan, which reports lipid packing based on local polarity and water penetration^33^, quantified as the Generalised Polarisation (GP) parameter^34^ (Extended Data Fig.4a). At 21°C, rods exhibited markedly higher GP values than the adjacent ER (Fig.3a), indicating tighter lipid packing and segregation into membrane domains of distinct order.

**Fig.3.**
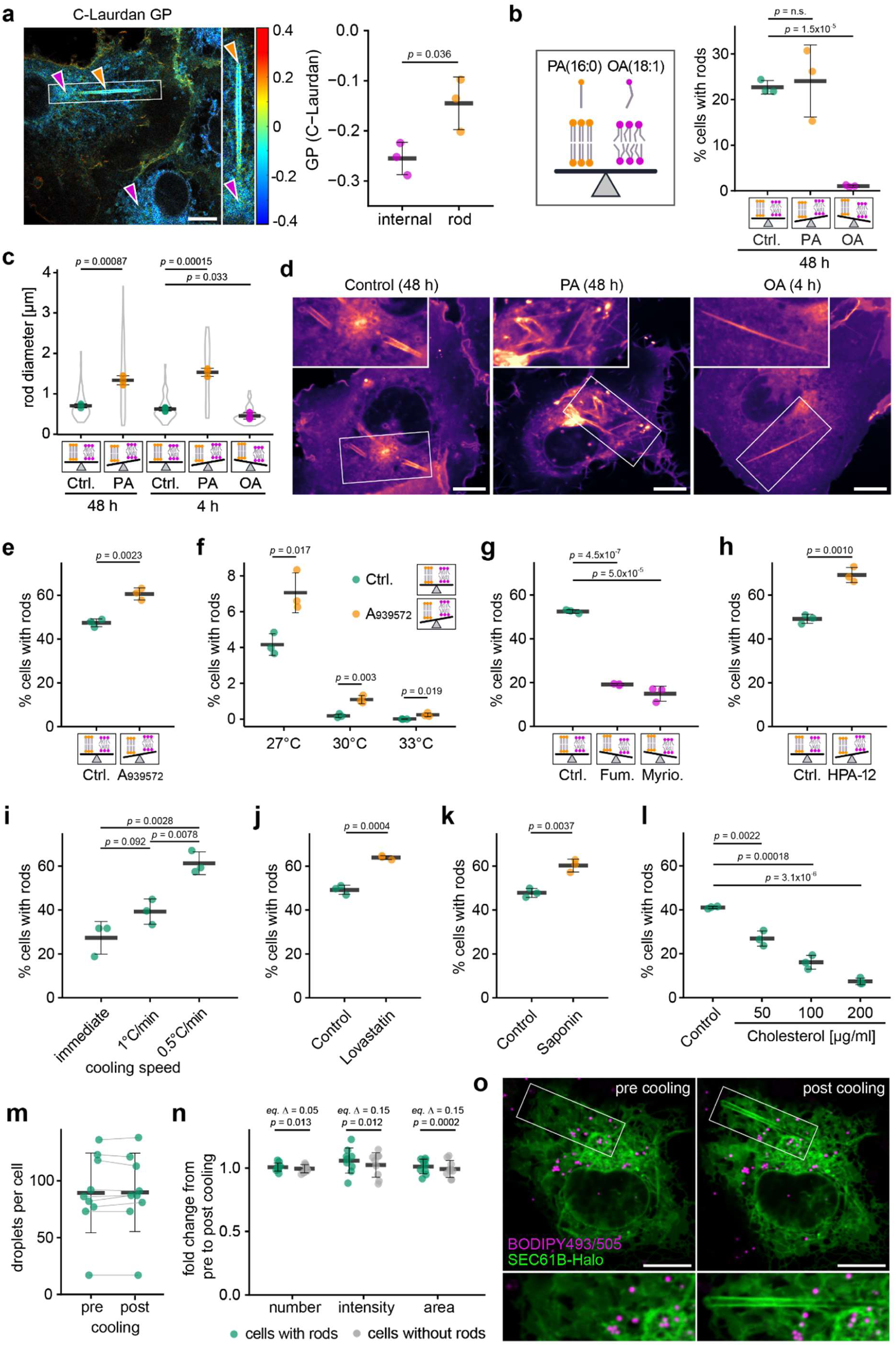
Rod formation is driven by lipid phase separation into solid-like domains. **a,** GP map of C-Laurdan-stained COS-7 cells, incubated at 21°C. Yellow arrowheads mark rods, magenta arrowheads internal membranes. GP values report lipid order, ranging from +1 (highly ordered) to -1 (disordered). Right: GP quantification for internal membranes, and rods. Black bar: mean of three independent experiments (≥5 cells per experiment); Error bars: SD. Scale bar: 10 µm **b,** Percentage of rod-positive cells expressing JF646-labelled SEC61B-Halo and JF-549-labelled SNAP-CDC42-HVR after 48h treatment with oleic acid (OA; 50 µM) or palmitic acid (PA; 50 µM) and cooling to 21°C. Black crossbars: mean of three independent experiments (coloured dots). Error bars: SD. Schematics illustrate shifts in lipid saturation. **c,** Rod diameters in cells from (**b**) after 48 h or 4 h treatment. Black crossbars: mean of three experiments; SD and violin plot distribution shown. **d,** Representative confocal sections of JF-549-labelled SNAP-CDC42-HVR cells showing PA- and OA-dependent rod morphology. Scale bar: 10 µm. **e-h,** Percentage of rod-positive CDC42-HVR–expressing cells treated with the indicated inhibitors before cooling to 21°C (or indicated temperatures in (**f**)) for 40 min. Black crossbars: mean of three experiments (coloured dots). SD shown. (**e,f**) A939572 (1 µM, 48 h); (**g**) fumonisin B1 (5 µM, 48 h) and myriocin (10 µM, 48 h); (**h**) HPA-12 (5 µM, 72 h). **i,** Live imaging of CDC42-HVR cells cooled from 37°C to 21°C using a CherryTemp device and different cooling protocols (immediate drop, 1°C/min ramp, or staged cooling: immediate drop to 28°C, followed by 0.5°C/min ramp). Black bar: mean of three independent experiments; Error bars: SD. **j,** Rod-positive cells after lovastatin treatment (2.5 µM, 72 h), quantified as in (**e-h**). **k,l,** Rod-positive cells after cholesterol extraction with saponin (30 µg/ml, 15 min) (**k**) or after loading with soluble cholesterol (30 min) (**l**), prior to cooling to 21°C for 40 min and quantified as in (**e-h**). **m,** Quantification of lipid droplets in JF646-labelled SEC61B-Halo-expressing cells stained with BODIPY493/503 before (37°C) and after cooling (21°C, 40 min). Nine paired cells quantified; black crossbar: mean; SD shown; grey lines link paired measurements. **n,** Fold changes (post/pre cooling) in lipid droplet number, intensity, and area from (**m**), grouped by cells with or without rods. Statistical equivalence is assessed by two one-sided test (TOST) with pre-defined equivalence margin Δ of 5% (number) and 15% (intensity and area). **o,** Representative confocal images corresponding to (**m**). Scale bars: 10 µm. Statistical significance was assessed by unpaired two-sided Student’s *t*-tests unless indicated otherwise.

To test whether lipid saturation contributes to rod formation, we manipulated cellular lipid composition by supplementing cells for 48h with either saturated palmitic acid (PA; 16:0) or monounsaturated oleic acid (OA; 18:1 cis-9). PA did not significantly change the number of rod-forming cells but produced strikingly enlarged and irregular rods, including incompletely wrapped, triangular profiles (Fig.3b-d). In contrast, OA completely abolished rod formation (Fig.3b). At shorter, non-steady-state incubation times, rods still forming in OA-treated cells had significantly reduced diameters (Fig.3c). Lipid droplet abundance was comparable under PA and OA treatment (Extended Data Fig.4b), excluding differential droplet induction as a confounding factor.

Pharmacological perturbation of lipid desaturation yielded similar results. Inhibition of stearoyl-CoA desaturase-1 (SCD1) with A939572^35^, enhanced rod formation at 21°C (Fig.3e) and raised the threshold temperature for their appearance, promoting formation at 27°C, 30°C, and even 33°C (Fig.3f). We next assessed the contribution of ceramides – highly saturated ER lipids with strong packing propensity^36,37^. Blocking ceramide synthesis via inhibitors of serine palmitoyltransferase (myriocin) or ceramide synthase (fumonisin B1) impaired rod formation (Fig.3g), whereas inhibition of ceramide export to the Golgi using the CERT transport protein inhibitor HPA-12 enhanced it (Fig.3h). These results implicate ceramides, and possibly related early sphingolipid intermediates, as contributors to rod formation. Across these experiments, shifting the balance toward saturated lipids promotes rod formation, and *vice versa* for unsaturated lipids, consistent with a lipid phase-separation mechanism.

Because lipid diffusion is rate limiting for membrane demixing, we tested whether rod formation depends on cooling kinetics. Rapid temperature drops across the 28–21 °C transition reduced the fraction of rod-containing cells (Fig.3i), consistent with elevated membrane viscosity constraining coalescence of saturated lipid domains. Conversely, prolonged incubation at lower temperature promoted rod formation: incubation at 16 °C for 3 h yielded more rod-containing cells than incubation at 21 °C for 40 min (Extended Data Fig.4c), as expected for a diffusion-controlled phase separation process.

The exceptional robustness (Supplementary Video9) and long-range order of rods suggest solid-like/gel phase characteristics. Cholesterol prevents such solidification by disrupting packing of saturated lipids. However, its concentration in the ER is low, potentially predisposing this compartment to solid-like phase formation. Further cholesterol depletion, either by inhibiting biosynthesis with lovastatin or by acute cholesterol extraction with saponin^38^ before cooling, markedly enhanced rod formation (Fig.3j,k). Post-formation treatment with digitonin progressively fragmented the surrounding ER but left rods intact (Extended Data Fig.4d), indicating membrane detergent resistance characteristic of high lipid order. Conversely, loading cells with cholesterol robustly suppressed rod formation (Fig.3l). Together, these findings indicate that rods comprise solid-like, compositionally distinct membrane domains that form within the cholesterol-poor ER. By sequestering saturated lipids, rods function as sinks for membrane-rigidifying components, thereby enriching the remaining ER in unsaturated lipids and increasing membrane fluidity. Because lipid droplets originate from the ER and can remain physically connected to it, we tested whether they contribute to this lipid redistribution. Quantification of lipid droplet number and volume before and after cooling in rod-forming and non-forming cells revealed no significant changes (Fig.3m-o), arguing against a role for lipid droplets in this process. We therefore conclude that rod formation represents a homeoviscous balancing mechanism in which sequestration of solidifying lipids into rods increases the fluidity of the remaining ER membranes.

Together, these findings demonstrate that rod formation depends on both temperature and lipid composition and is driven by lipid phase separation. Saturated lipids, including ceramides, coalesce into large, highly ordered, solid-like domains that segregate into rods, while the remaining ER retains a predominantly fluid, unsaturated character.

### Lipid phase separation segregates ER-tubulating proteins to promote membrane flattening

How does lipid demixing drive rod formation? Previous studies have shown that ER proteins, including those that shape ER morphology, can occupy distinct microdomains within the membrane^2–4^. We therefore investigated how ER proteins distribute within and outside rods.

A broad panel of proteins containing single-pass transmembrane domains (TMD; SEC61B, VAPA, the Calnexin TMD, CYTB5 and LYRIC) robustly localised to rods (Fig.4a left and Extended Data Fig.5a,b). Membrane topology was critical: cytosolically tagged VAPA or Calnexin-TMD entered rods (Fig.4a left), whereas luminally tagged versions were excluded (Fig. 4a right), consistent with the exclusion of luminal KDEL and Calreticulin due to multilamellar packing (see Fig.2g,h). Also, the multi-pass proteins AMFR (GP78; six-pass) and CEPT1 (ten-pass) partitioned into rod membranes (Extended Data Fig.5a,b), together indicating that proteins with conventional ER-spanning helices and lacking bulky luminal domains can incorporate into these structures.

**Fig.4.**
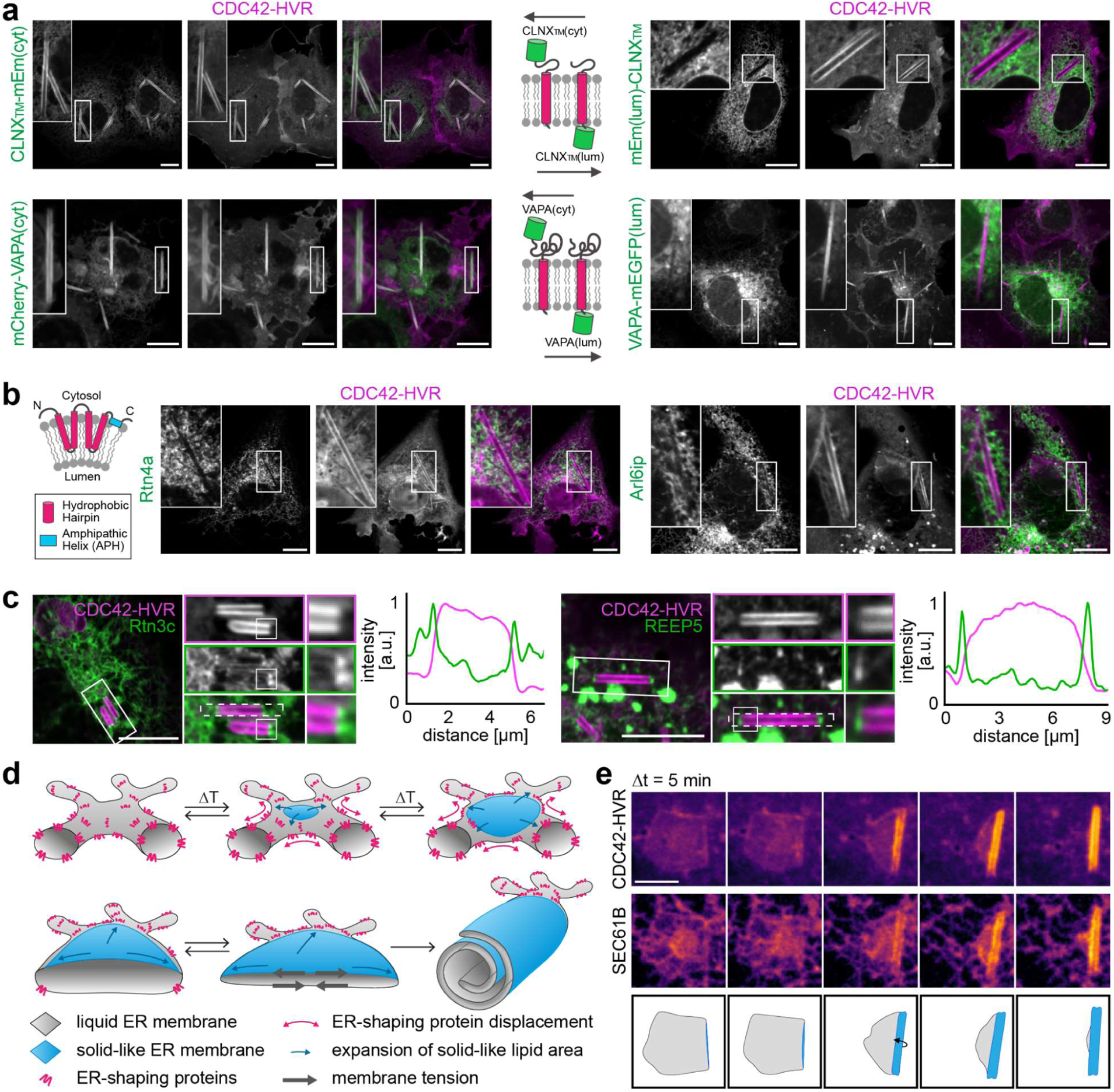
Exclusion of ER-tubulating proteins from rods. **a-c,** Confocal micrographs of cells expressing the indicated ER marker constructs (**a**) or reticulon homology domain (RHD)-containing proteins (**b,c**). Cells were cooled to 21°C for 40 min prior to imaging. Localisation on rods versus exclusion form rods: CLNX_TM_-mEm(cyt) (30/0), mEm(lum)-CLNX_TM_ (3/32), mCherry-VAPA(cyt) (21/0), VAPA-GFP(lum) (0/20), Rtn4a (0/31), Arl6ip (0/27), Rtn3c (0/26), REEP5 (0/28). (**a**, middle): fluorescent tag orientation relative to ER membrane. (**b**, left): Membrane binding topology of RHD-containing proteins. **c,** Enrichment of Rtn3c and REEP5 at end-to-end contact sites between ER tubules and rods. Integrated fluorescence intensity within dashed regions is quantified. **d,** Model of temperature-induced rod formation. Top: Cooling drives lipid phase separation in the ER, expanding solid-like membrane domains (blue arrows) that exclude proteins incompatible with rigidified bilayers. Bottom: Reticulon segregation (magenta symbols/arrows) permits progressive membrane flattening. Asymmetric solid-like domain distribution across ER bilayers generates membrane tension (grey arrows), coupled to ER lumen volume reduction, driving membrane wrapping into multilamellar rods. **e,** Time-lapse images of cells expressing CDC42-HVR and SEC61B during cooling to 21°C. Bottom: Schematic segmentation showing rolling of an ER sheet (grey) into a rod (blue). Scale bars: 10 µm (**a-c**); 5 µm (**e**).

In contrast, reticulon family proteins, key mediators of ER tubulation, were conspicuously excluded from rods (Fig.4b,c). Reticulons use their reticulon homology domains (RHDs) to insert paired hydrophobic hairpins and amphipathic helices (APHs) into the cytosolic leaflet to induce curvature. REEP5 (DP1) and ARL6IP1, other RHD-containing, curvature-promoting proteins^12,39^ were similarly excluded. The RHD proteins accumulated at end-to-end contacts between ER tubules and rods, suggesting that ordered rod membranes are incompatible with RHD insertion (Fig.4c and Supplementary Video10). Deletion of the APH in REEP5 or Rtn4 did not restore rod localisation (Extended Data Fig.5c), nor did mutations in REEP5 predicted to impair oligomerisation (analogous to hereditary spastic paraplegia type 31 mutations in REEP1^40^). Thus, exclusion of RHD proteins is primarily driven by steric incompatibility between their wedge-like hairpins and the densely packed rod membranes, independent of their oligomerisation-based scaffolding function or the APH^12–14^.

Together, these findings support a model in which lipid phase separation excludes ER-tubule promoting proteins, allowing flat, ordered membrane domains to expand. Stochastic, asymmetric growth of coexisting lipid phases across the two ER bilayers then promotes the wrapping of planar sheets into continuous multilamellar rods (Fig.4d). Consistently, live imaging captured ER sheets transforming into rods (Fig.4e and Supplementary Video11), and occasionally rod structures connected to giant membrane sheets with sharp, linear edges, possibly representing stalled intermediates *en route* to fully wrapped multilamellar rods (Extended Data Fig.5d and Supplementary Video12).

This model predicts that rods still retain fluid membrane territories - necessarily along highly curved edges where enrichment in loosely packed, unsaturated lipids is required - and within alternating apposed bilayers of the multilamellar structure. Accordingly, fluorescence recovery after photobleaching (FRAP) of the single-pass ER protein SEC61B revealed a reduced but detectable mobile fraction within rods (Extended Data Fig.5e), indicating the presence of a diffusive, yet restricted phase in rods.

### Rods form at physiological temperatures and *in vivo* in lung cells

To explore whether rod structures can form *in vivo* and under physiological conditions, we investigated cells naturally enriched in saturated fatty acids. Alveolar Type II (AT-2) lung cells are a prime example, as they contain organelles known as lamellar bodies (LBs) that consist of up to 40% of the saturated lipid dipalmitoylphosphatidylcholine (DPPC). AT-2 cells store pulmonary surfactant in LBs and secrete it into the alveolar space^41–43^. Like rods, LBs exhibit a multilamellar organisation, despite being spherical, discontinuous and structurally dependent on surfactant proteins. The high DPPC content is thought to confer mechanical stability to the multilayered membranes and facilitate structural remodelling of surfactant during respiration^44^.

We first analysed Kras^LA2^ mouse lung tumours, which are strongly enriched in AT-2 cells. Electron microscopy and volume EM of tumour AT-2 cells isolated and fixed at 21°C revealed rod structures that could be fully reconstructed in 3D (Fig.5a-d and Supplementary Video13). These rods were indistinguishable in shape, diameter and length from those seen in COS-7 cells, yet clearly distinct from the canonical LBs present in the same cells. In many instances, dark osmophilic enrichments were recorded at rod termini (Fig.5e), consistent with the previously observed local accumulation of curvature-promoting proteins (Fig4b,c). Importantly, rods also formed in samples processed entirely at 37°C, with identical morphology to those formed at lower temperatures (Fig.5f), albeit at a lower frequency. AT-2 cells in lungs derived from wild-type C57BL/6 mice displayed comparable rod formation (Fig.5g,h), indicating that their occurrence is not tumour-specific. Consistent with the temperature-dependence of rod formation, rod abundance increased as the processing temperature decreased (Fig.5h). Detailed ultrastructural analysis further highlighted the exceptional rigidity of rods, which could deform neighbouring organelles and enclose entire mitochondria (Fig.5i; Extended Data Fig.6b). It also revealed intermediate morphologies consistent with sheet-to-rod transition events and the continuity of rods and regular ER (Fig.5j, Extended Data Fig.6b,c and Supplementary Videos14 and 15). Together, these findings establish the physiological existence of rods at body temperature when the membrane lipid composition is tuned toward high saturation, highlighting their dual dependence on environmental and metabolic factors.

**Fig.5.**
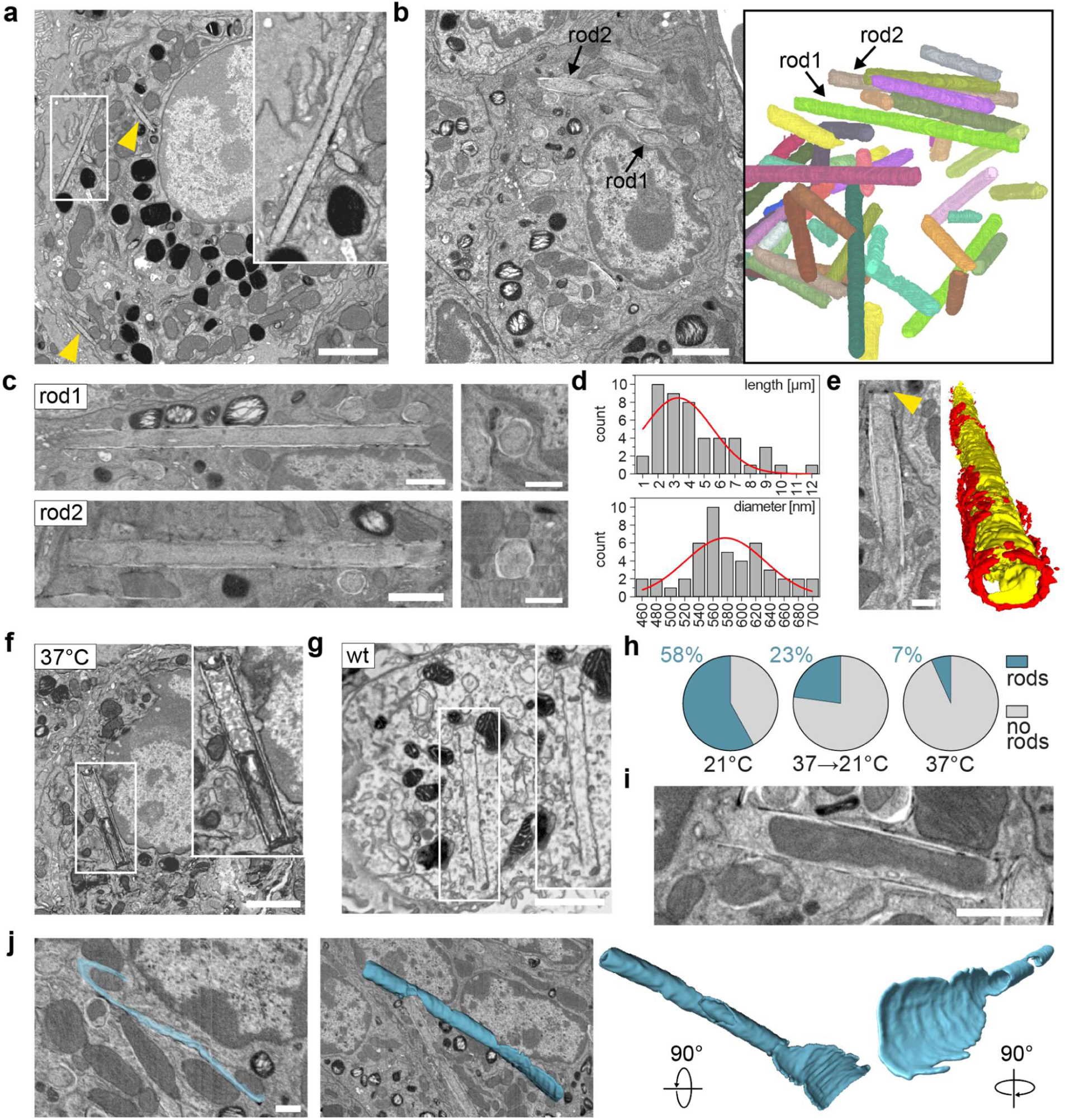
Rods form in mouse lung AT-2 cells at physiological temperatures. **a,** Array tomography SEM image of an AT-2 cell from a *Kras^LA^*^2^ lung tumour fixed at room temperature, showing rods (arrowheads and inset). **b,** FIB-SEM analysis of a *Kras^LA^*^2^ lung tumour fixed at room temperature. Left: 2D slice from the 3D volume. Right: 3D reconstruction of segmented rods. **c,** Rods from (**b**) sectioned along the long (left) and short axis (right). **d,** Distribution of rod diameters (top) and lengths (bottom). **e,** Osmophilic enrichment at rod terminus, visible as two dark foci in cross-section (left, arrowhead) and segmented in red in a 3D reconstruction (right; rod body in yellow). **f,** Array tomography image of an AT-2 cell from *Kras^LA^*^2^ lung tumour fixed at 37C. **g,** Array tomography image of an AT-2 cell from wild-type C57BL/6 mouse lung fixed at room temperature. **h,** Quantitation of wild-type lung AT-2 cells with (blue) or without rods (grey) after fixation at 21°C (n=124), 37°C (n=119), or upon cooling from 37°C to 21°C for 20 min (n=105). **i,** EM image showing rod enclosing and deforming mitochondria. **j,** Putative sheet-to-rod intermediate from FIB-SEM reconstruction (segmented in blue). Scale bars: 2 µm (**a,b,f,g,i**), 1 µm (**c,** left), 500 nm (**c,** right**, e, j**).

## Discussion

We have structurally and mechanistically characterised a previously unrecognised manifestation of organelle polymorphism: the temperature- and lipid saturation-dependent formation of giant, rigid membrane rods emerging from the ER network. Given their extraordinary dimensions and likely unmatched persistence length in cell-biological systems, it is remarkable that rods have escaped prior characterisation. Nonetheless, earlier observations are consistent with their existence, including ‘bar-like’ EM structures in AT2 cells^45,46^ and ‘long, thin solid-like structures’ reported in *AdipoR1/2* deficient cells with impaired fatty acid desaturation following palmitate treatment^47^.

Rod formation represents a striking mode of endomembrane phase behaviour: saturated lipids segregate into stable, micrometre-scale solid-like domains that confer long-range orientational order to the bilayer. These domains are biophysically distinct from the transient liquid-ordered “rafts” of the plasma membrane. In mammalian cells, solidification at this scale has been viewed as both detrimental and unlikely, as reduced fluidity compromises membrane plasticity and protein function. Accordingly, cells prevent this transition through lipid unsaturation and cholesterol buffering. Consistent with this, cholesterol extraction induces solid-like regions in the plasma membrane^26^. However, ER membranes contain only a small fraction of cholesterol compared to the plasma membrane (∼5% versus ∼30-40%)^17^, creating conditions permissive for demixing. We show that modulating cholesterol levels alters the propensity for rod formation in the ER. Solid-like endomembrane phase behaviour has been reported earlier in the yeast nuclear envelope and bud necks^20,21^ and, under extreme conditions, in mammalian cells loaded with saturated fatty acids or exposed to hypotonic swelling and cooling^24,25^. Here, we provide the first direct visualisation of reversible micrometre-scale lipid phase separation in living mammalian cells, establishing a tractable model for investigating solid-like phase transitions in an otherwise unperturbed cellular context.

We further show that lipid demixing drives the segregation of proteins into discrete endomembrane territories, providing a mechanistic basis for the spatial organisation of the ER. Solid-like lipid domains in rods exclude proteins containing reticulon homology domains, which require lipid-packing defects for membrane insertion and curvature generation^48^. Although microscale rods dissolve above the transition temperature, nanoscale solid-like domains may persist at physiological temperature. This behaviour is consistent with membrane dynamics close to criticality, where compositional fluctuations are sustained and domain size and lifetime are not fixed^49^. Similar solid-like nanodomains have been reported in cholesterol-depleted plasma membranes^50^. Such lipid phase separation may underlie the previously reported compartmentalisation of ER proteins, including membrane-shaping factors, and may act alongside protein-based mechanisms such as restricted diffusion or scaffolding^3^. Changes in lipid metabolism may therefore directly tune ER architecture and shift the balance between sheets and tubules. Such morphological and spatial modulation provides a route by which environmental cues, stress, or cell-cycle state could rapidly reprogram ER functions in biosynthesis, signalling and inter-organelle communication^7,23^. Future work will explore ordered ER nanodomains in living cells and determine how lipid-driven segregation is integrated with protein-mediated organization. Our findings highlight membrane fluidity as a key determinant of ER architectural plasticity: Reticulon-mediated tubulation requires a permissive lipid environment low in saturated lipids and cholesterol – a constraint that likely contributes to the compositional gradient from the fluid ER to the more ordered peripheral membranes.

Rods exhibit a continuous multilamellar architecture that appears unprecedented among mammalian organelles. This multilamellarity implies membrane properties that overcome electrostatic and steric repulsion between closely apposed leaflets. Comparable multilamellar organisations are rare and discontinuous, occurring only in specialised contexts such as MHCII processing lysosomes^51^, lamellar bodies of alveolar type II (AT-2) cells used in this study^42^ and ER-derived replication organelles formed during positive-strand RNA virus infection^52^. ER rods thus represent a yet undescribed multilamellar membrane organisation, providing a model to probe the biophysical determinants of multilamellar membrane formation and stability.

Methodologically, maintaining physiological temperature should become standard practice for EM sample preparation when studying membrane organelle morphology in tissue-derived samples. We observed rod formation at 37°C in AT-2 cells enriched in saturated lipids, suggesting that tissues with elevated SFA/UFA ratio support rod formation at physiological temperature. Metabolic rewiring in cancer or during viral infection may similarly promote rod formation. Whether the *in vivo* demixing temperature exceeds that observed in cultured cells remains unresolved, particularly given that culture lipidomes are shaped by bovine serum, which contains roughly twice the monounsaturated and half the polyunsaturated lipid species present in native tissue^53^.

Finally, sequestration of solidifying lipids into rods enriches the surrounding ER in unsaturated species, increasing bulk membrane fluidity. Rod formation may therefore function as a homeoviscous buffering mechanism that limits ER stiffening during acute thermal or metabolic stress while preserving reticulon-dependent tubulation. By stabilising ER architecture, this process may modulate stress signalling triggered by membrane rigidification. Defining how lipid phase separation couples to ER plasticity and cellular fitness will be essential for understanding how the ER maintains structural and functional homeostasis in physiological and pathological states.

## Methods

### Plasmids

mCitrine(YFP)-CDC42-HVR (EPPEPKKSRRCVLL, this study), mCherry-CDC42-HVR (this study), SNAP-CDC42-HVR (this study), mCherry-RHOA-HVR (QARRGKKKSGCLVL, this study), mCherry-RAC1-HVR (CPPPVKKRKRKCLLL, this study), YFP-KRAS-HVR (KKKKKKSKTKCVIM, this study), GalT-YFP, CFP-Rab5, YFP-RAB7, SEC61B-mEm (mEmerald GFP, this study), SEC61B-Halo (this study), PTP1B-CFP (gift of P. Bastiaens Lab), CytERM-DsRed2 (addgene #55835), CytERM-mGFP (addgene #62237), mEm-KDEL, mEm(lum)-CLNX-TM (addgene #186961), CLNX-TM-mEm(cyt) (addgene #186962), mCherry-Cyt5B (addgene #182579), GFP-RTN4A (addgene #86684), Rtn3c-mEm (addgene #186963), REEP5-mEm (addgene #186964), ARL6IP-mEm (addgene #186965), Calreticulin-mEm (addgene #54023), mCherry-VAPA (addgene #226407), VAPA-GFP (this study), mCherry-CEPT1 (addgene #119079), AMFR(gp78)-GFP (addgene #37375), Lyric-mCherry (addgene #226563), GFP-RTN4HD (addgene #86686), GFP-RTN4-RHD-ΔAPH (this study), mScarlet-REEP5-ΔAPH (this study), REEP5-PWmut-mEm (P63L/W84R) (this study)

### Cell culture and transfection

COS-7 (ATCC CRL-1651), HEK293T (ATCC CRL-3216), HeLa (ATCC CRL-2), MDCK II (ATCC CRL-2936), U-2 OS (ATCC HTB-96) and C2C12 cells (ATCC CRL-1772) were maintained in Dulbeccośs modified Eagle’s medium (DMEM, Thermo Fisher) supplemented with 10% fetal bovine serum (FBS, PAN Biotech) and penicillin-streptomycin (100 U/ml, Thermo Fisher) at 37°C and 5% CO₂. Cells were grown to ∼60-80% confluency and transfected with Lipofectamine 3000 (Thermo Fisher) following the manufacturer’s protocol, except MDCK II cells, which were transfected with Effectene (Qiagen). Cells were incubated overnight before experiments. For live-cell imaging, cells were plated on 8-well glass-bottom chamber slides (Ibidi) or 35 mm glass bottom dishes (MattTek). For fixed-cell experiments, cells were seeded on 13 mm round glass coverslips (Karl Hecht) placed in 12-well tissue culture plates (Sigma Aldrich).

### Labelling JF dyes with HaloTag and SNAPTag ligand

Cells expressing either HaloTag or SNAPTag were incubated in complete media with 1 µM JF646 HaloTag ligand or 500 nM JF549 SNAPTag ligand at 37 °C for at least 30 min, washed twice and equilibrated at 37 °C for 30 min before experimentation.

### Cell fixation and immunofluorescence staining

Cells were washed once with pre-warmed PBS containing calcium and magnesium (PBS+/+, PAN-Biotech), equilibrated to the same temperature as the culture conditions. To preserve rod structures, fixation was performed for 10 min using 4% paraformaldehyde (Merck) supplemented with 0.1% glutaraldehyde (EM grade, Electron Microscopy Sciences) at the corresponding temperature, followed by washing three times with PBS (10 min per wash).

For immunostaining, cells were permeabilised with 0.1% Triton X-100 (Sigma Aldrich) and 100 mM Glycine in PBS, followed by three washes with PBS. Blocking was performed using 3% BSA (Cell Signaling) in PBS for 30 min. Cells were incubated overnight at 4°C with mouse monoclonal anti-α-tubulin antibody (Thermo Fisher, A11126; 1:1,000 in blocking buffer), then washed three times with PBS for 10 minutes. Secondary staining was performed by incubating for 1 h in the dark with an Alexa Fluor647 goat anti-mouse secondary antibody (Thermo Fisher, A21237; 1:500), combined with Alexa568-conjugated phalloidin (Thermo Fisher, 1:500) in blocking buffer. Cells were again washed three times with PBS for 10 min. For mounting, coverslips with fixed cells were inverted onto glass slides using ProLong Gold Antifade Mountant (Thermo Fisher).

### Temperature dependent induction of rod formation

Unless otherwise specified, for live-cell imaging of rod formation the culture medium was replaced by FluoroBrite DMEM supplemented with 1% FBS. Rod formation was induced by transferring cells from 37°C to an air-conditioned microscope environment at 21°C and incubating them for 40 min to allow gradual cooling and thermal equilibration. For experiments involving subsequent staining or washing steps, all solutions were pre-equilibrated to 21°C to preserve rod integrity. For standard imaging experiments, the microscope stage was maintained at 21°C; where necessary, ice blocks were placed in the incubation chamber to achieve and stabilise this temperature, which was continuously monitored using a cable-probe thermometer.

For the analysis of the critical temperature for rod formation, the culture medium was replaced by FluoroBrite DMEM containing 1% FBS, and cells were transferred from 37°C to a cooling incubator preset to different temperatures between 11°C and 30°C (Binder). Cells were placed on an equilibrated aluminium metal plate (20 x 20 x 1 cm) covered with a thin water film for rapid heat transfer. Cells were kept at the target temperature for 40 minutes before fixation. Z-stacks were acquired from 40 fields of view and the number of rod-containing cells was quantified.

For live cell imaging experiments involving precise and dynamic temperature control, a CherryTemp microscope on-stage device was used (Cherry Biotech). In brief, cells seeded in an adhesive silicone culture-insert (ibidi) on 24×60 mm glass coverslips were mounted in a closed chamber comprising a 0.5 mm thick PDMS spacer and a thermalisation chip positioned on top according to the manufacturer’s instructions.

To assess the effect of temperature on demixing kinetics, the CherryTemp device was used with three cooling protocols: 1) an immediate temperature drop from 37 to 21°C, followed by a 40 min hold; 2) slow cooling from 37 to 21°C at 1°C/min followed by a hold, with a total duration of 40 min; and 3) immediate drop from 37 to 28°C followed by slow cooling from 28 to 21°C at 0.5°C/min and a hold, with a total duration of 40 min. Immediately thereafter, z-stacks from 40 randomly selected fields of view were acquired, and rod-containing cells were quantified.

### Spinning disk confocal imaging

Images and movies were acquired on either a Nikon CSU-W spinning disk microscope equipped with a 100x Oil CFI Plan Apo λ/NA 1.45/WD 0.13 objective and an Andor iXON DU-888 EMCCD camera, or on a Nikon CSU-W1 SoRa spinning disk microscope equipped with a 100x Oil Plan Apo λ DIC N2/ NA 1.45/WD 0.13 objective and a Hamamatsu ORCA-Fusion digital sCMOS camera. The following excitation (Ex) and emission (Em) settings were used: CFP Ex: 445 nm laser, Em: 472/30 nm; mEmerald-GFP, BODIPY 493/503, mCitrine-YFP Ex: 488 nm laser, Em: 525/50 nm; mCherry, mScarlet, ERTracker Red, MitoTracker Red, JF549-SNAP Tag ligand Ex: 561 nm laser; Em: 600/52 nm; JF646-Halo Tag ligand Ex: 638 nm laser, Em: 708/75 nm.

Image stacks were deconvolved using Nikon NIS-Elements AR software (version 5.42.07, Nikon Instruments) with the built-in 3D deconvolution module implementing the Richardson-Lucy method.

### BODIPY 493/503 staining

A 1 mM BODIPY 493/503 (Sigma-Aldrich) stock in DMSO was diluted in FluoroBrite DMEM imaging medium (ThermoFisher) to 1 µg/mL. For *post–temperature drop* staining, cells were incubated at 21°C for 40 min to induce rod formation, washed with FluoroBrite pre-cooled to 21°C, stained with the 21°C BODIPY working solution for 40 min at 21°C, and imaged immediately. For *pre-temperature drop* staining, cells were washed once with DMEM warmed to 37°C, stained with the BODIPY working solution for 30 min at 37°C, washed with FluoroBrite warmed to 37°C, shifted to 21°C for 40 min to induce rod formation, and imaged immediately. For lipid droplet quantification, cells were stained with the BODIPY working solution for 30 min at 37°C, imaged, then incubated at 21°C for 40 min to induce rod formation and image again.

### ERTracker and MitoTracker staining

Cells were cooled to 21°C for 40 min to induce rod formation, washed with FluoroBrite pre-cooled to 21°C and stained for 45 min with 1 μM ERTracker-Red or 500nM MitoTracker-Red (Thermo Fisher) in FluoroBrite at 21°C. Cells were washed twice with 21°C FluoroBrite and imaged.

### ATP Depletion

Cells were incubated with PBS containing 1 mM CaCl₂ and 1 mM MgCl₂ supplemented with 20 mM 2-deoxy-D-glucose and 8 mM sodium azide (Carl Roth) for 15 min at 37°C, followed by an additional 40 min at 21°C to induce rod formation, and subsequently fixed.

### Interference with cytoskeletal dynamics

Cells were incubated at 37°C for 30 min with 10 μM Cytochalasin D or 15 µM Nocodazole, or with 25 µM Y-27632 for 10 min (all Sigma Aldrich), cooled to 21°C for 40 min to induce rod formation, fixed and subsequently imaged.

### Cell Supplementation with palmitic acid and oleic acid

Palmitic acid (PA) and oleic acid (OA) were complexed with fatty acid-free BSA at a 2:1 ratio (PA:BSA or OA:BSA 2 mM:1 mM), BSA alone was used as control. Cells were incubated with PA:BSA or OA:BSA at 100 µM:50 µM or with BSA alone at 50 µM in complete medium for 4 h or 48 h. After 24 h medium was replaced with freshly diluted PA:BSA or OA:BSA. Cells were cooled to 21°C for 40 min to induce rod formation prior to imaging.

### Inhibitor treatments

Cells on 13-mm glass coverslips were treated with the inhibitors A939572 (Sigma Aldrich), myriocin (Sigma Aldrich), fumonisin B1 (Stem Cell Technologies), HPA-12 (Sigma Aldrich) and lovastatin (Thermo Fisher) at the indicated concentrations and for the specified durations. After treatment, cells were cooled to 21 °C for 40 min to induce rod formation, then fixed and the number of rod-containing cells quantified.

### Cholesterol extraction and supplementation

Cells grown on 13-mm glass coverslips were incubated in DMEM without FBS for 1 h, treated with 30 µg/ml saponin (Carl Roth) for 15 min at 37°C, and then cooled to 21°C for 40 min to induce rod formation. Cells were subsequently fixed and the fraction of rod-containing cells was quantified. For cholesterol loading, cells were incubated in DMEM without FBS for 1 h, treated with water-soluble cholesterol (C4951, Sigma Aldrich) for 30 min at 37°C, cooled to 21°C for 40 min, fixed, and quantified. For live-cell imaging of digitonin treatment, cells were maintained in FluoroBrite DMEM containing 1% FBS at 21°C and imaged during exposure to 10 µM digitonin (Cayman Chemicals).

### Transmission electron microscopy

Cells were either chemically fixed or cryo-immobilised by high pressure freezing. Chemical fixation was performed with 2% (w/v) formaldehyde and 2.5% (v/v) glutaraldehyde in 0.1M phosphate buffer. Samples were post-fixed with 1% (w/v) tannic acid, 1% (v/v) osmium tetroxide and 0.5% (w/v) uranyl acetate, dehydrated through a graded ethanol series, and embedded in the PolyBed® 812 resin (Polysciences).

High pressure freezing (Leica EM ICE) of cell monolayers on sapphire discs was followed by freeze substitution (Leica AFS2) in 1% (w/v) osmium tetroxide, 1% (v/v) glutaraldehyde and 1% water in acetone for 36 h at -90°C, 8 h from -90°C to -50°C, for 6 h at -50°C to -20°C, for 12 h at -20°C and 3 h at -20°C to +20°C. Contrast was enhanced by incubation in 0.1% (w/v) uranyl acetate in acetone before resin infiltration with PolyBed® 812 resin (Polysciences).

Ultrathin sections (60 - 80 nm) were cut and stained with lead citrate before imaging (Leica Microsystems). For correlative confocal light - and transmission electron microscopy (CLEM) fluorescent landmarks and gridded dishes (ibidi) were used for targeted ultramicrotomy. Electron micrographs of thin resin sections were correlated to optical slices from the confocal z-stack using cellular landmarks and ER/nuclear fluorescence signals with the ecCLEM plugin in Icy^54^.

For immunolabeling, cells were fixed for 2 h at room temperature with 2% (w/v) formaldehyde and 0.5 % (v/v) glutaraldehyde in 0.1 M phosphate buffer and processed as described in^55^. Briefly, samples were infiltrated with 2.3 M sucrose, frozen in liquid nitrogen, and sectioned at cryo-temperatures. Sections were blocked with PBS containing 1% BSA and 0.1% glycine, labelled with rabbit polyclonal anti-GFP (1:200, abcam 6556, Abcam) and 12-nm colloidal gold (Dianova), and contrasted with 3% (w/v) tungstosilicic acid hydrate in 2.8% polyvinyl alcohol (w/v)^56^.

All sections were examined at 80 kV with a Zeiss EM 910 microscope using a Quemesa CDD camera and iTEM software (Emsis).

### Electron tomography

Cultured cells were either chemically fixed or cryo-immobilised by HPF and embedded as described in the previous paragraph. Serial sections with a thickness of 300 nm were obtained with a Leica UC6 ultramicrotome (Leica Microsystems) and mounted on finder grids. After post-contrasting with uranyl acetate and lead citrate the samples were imaged with an F30 electron microscope (Thermo Fisher). Tilt series ranging from -63° to 63° were acquired followed by reconstruction with the IMOD software package^57^. Segmentation and visualization were done with Microscopy Image Browser^58^ and ORS Dragonfly (https://dragonfly.comet.tech/). Resin-embedded mouse lung tumour sections (See Preparation of mouse lung tissues for SEM for sample preparation; 250 nm thick) were acquired using a Leica ARTOS ultramicrotome and collected onto glow-discharged formvar-coated copper slot grids (2 x 1 mm). Tilt series of alveolar Type II (AT-2) cells containing rods were acquired using a 120 kV Talos L120C TEM (Thermo Fisher Scientific) equipped with a model 2020 advanced tomography holder (Fischione). Data acquisition utilised SerialEM software^59^ at a magnification corresponding to either 0.9663 or 0.7573 nm pixels, with 1° tilt increments across a ±60° angular range. Tomograms were reconstructed using Etomo (part of the IMOD software package, Ver. 4.11^60^), employing patch tracking and weighted back-projection with the simultaneous iterative reconstruction-like filter. Final tomogram movies were generated in Quicktime.

### C-Laurdan spectral imaging

C-Laurdan imaging was performed as previously described^34,61^. Briefly, cells were washed 2x with PBS, and incubated with 10 µg/mL C-Laurdan (Tocris Bioscience) for 20 min at RT. Finally, the cells were washed with PBS and then imaged at room temperature with a 63X water immersion objective on a Leica SP8 confocal microscope with spectral imaging with excitation at 405 nm. The emission was collected as two images: 420–460 nm and 470–510 nm. MATLAB (MathWorks, Natick) was used to calculate the two-dimensional (2D) GP map, where GP for each pixel was calculated as previously described^62^. Briefly, each image was background subtracted and thresholded to keep only pixels with intensities greater than 3 standard deviations of the background value in both channels. The GP image was calculated for each pixel with the following equation (where G is the G-factor and I is intensity at wavelength):

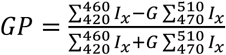

The G-factor was determined before each experiment using a previously published protocol^34^. GP maps (pixels represented by GP value rather than intensity) were imported into ImageJ. The average GP of the internal membranes was calculated by determining the average GP of all pixels in a mask drawn on each cell just inside of the PM, excluding the rods. Regions of interest were drawn along the rods, and the average value reported. 7-20 cells were imaged and quantified per experiment.

### Quantification and statistical analysis

All quantified experiments were performed as independent triplicates unless otherwise specified. Data are represented as the mean ±s.d. Statistical significance was determined using unpaired, two-sided Student’s t-tests unless otherwise indicated. P-values for group comparisons are shown in the figures. For Fig.3n, statistical equivalence was assessed using the two one-sided test (TOST) procedure with predefined equivalence margin (Δ) of 5% for lipid droplet number and 15% for lipid droplet intensity and area. The equivalence margins were estimated based on the s.d. of the “cells without rods” group.

### Preparation of mouse lung tissues for SEM

Mice were humanely euthanised using carbon dioxide asphyxiation in accordance with the American Veterinary Medical Association (AVMA) Guidelines. All procedures were approved by the Institutional Animal Care and Use Committee (IACUC). After euthanasia, mouse lungs were excised. In the K-ras^LA2^ mice, visible lung tumours were dissected and cut into smaller pieces. In the C57BL/6 mice, the lungs were inflated with tyrode buffer at either 21°C or 37°C, then cut in smaller pieces (∼1 mm^3^). Some lung tissue pieces were incubated in tyrode buffer at 21°C or 37°C for 40 minutes before fixation. All tissue samples were then fixed with Karnovsky’s fixative (4% formaldehyde and 2% glutaraldehyde in 0.1 M cacodylate buffer) at either 21°C or 37°C before transport, depending on the experimental group. Samples were post-fixed in 2% osmium tetroxide and 1.5% potassium ferricyanide in 0.1 M sodium cacodylate for 1 hr at RT, washed with ultrapure water, and stained overnight with 1% aqueous uranyl acetate at 4°C. After additional water washes, the samples were treated with lead aspartate (0.066 g lead nitrate dissolved in 10 mL 30 mM aspartic acid; pH 5.5) at 65°C for 30 minutes and washed again. Tissues were dehydrated in graded ethanol (35%, 50%, 70%, 95%, and 100% x 5; 10 min. each) and then in propylene oxide (PO; 100% x 5; 10 min. each). The tissues were infltrated with increasing amounts of Polybed 812 resin (hard formulation) with PO (resin: PO; 1:3, 1:2, 3:1, and 100%) and finally embedded in 100% degassed resin and incubated at 65°C for 48 hrs. (EMS).

### Array tomography SEM workflow and quantification of AT-2 cells with rods in mouse lung and tumours

After polymerisation, resin-embedded mouse lung and tumour tissue blocks were trimmed with a Leica EM TRIM2 milling system. The block faces were then sectioned at 100 nm with a 45° ultra-diamond knife (Diatome) on a Leica ARTOS ultramicrotome. Sections were collected with a loop and placed on glow-discharged ITO coverslips, which were subsequently mounted onto 4-inch type-p silicon wafers with copper tape and placed on a 4-inch stage-decel holder in a GeminiSEM 450 (Zeiss). Images were collected using a four-quadrant backscatter detector (aBSD, Zeiss) with the electron beam operated at 3.0 kV with a 1.0 kV beam deceleration and 800 pA probe current. Low-resolution overview scans (150 nm pixel size) of individual resin sections were acquired using ATLAS 5 Array Tomography software (Fibics). High-resolution image mosaics (10 nm pixels; ∼60,000 x 35,000 pixels) were generated by stitching multiple fields, using automated stage movement. These mosaics were stitched and aligned computationally in ATLAS 5 to produce large-area images.

The stitched mosaics were imported into napari^63^, and the empanada plugin ver. 1.2^64^ was used to create 16,000 x 16,000 pixel image tiles (roughly 8 – 12 tiles per mosaic). Using the napari points layer function, AT-2 cells were annotated on each tile. AT-2 cells, distinguished by their characteristic lamellar body organelles^65,66^, were marked, as were AT-2 cells containing rods. Annotation layers were exported as CSV files and cell counts per image tile were aggregated by condition to give the total number of AT-2 cells with and without rods. For the mouse lung tissue, entire images were analysed (21°C: n = 124; 37°C: n = 119). For mouse lung tumour tissue, only two image tiles (16,000 x 16,000 pixels each) were quantified per condition, as AT-2 cells are the predominant cell type in these densely packed tumours (21°C: n = 300, 37°C: n = 379). Cell counts for AT-2 cells with and without rods were then plotted using GraphPad Prism (Ver. 10.5.0 for Mac).

### Mouse lung AT-2 cell FIB-SEM, rod segmentation, and analysis

Resin-embedded mouse tissue samples were mounted onto SEM stubs, silver painted, and carbon coated (10 nm) before insertion into the Zeiss Crossbeam 550 microscope. Target regions were located using SEM imaging at 1.5 kV accelerating voltage and 1.1 nA beam current. The FIB sample preparation workflow was carried out using the ATLAS3D (Fibics) sample preparation workflow, largely described in^67^. A 1 um platinum pad was deposited over the target cell, followed by milling of the tracking and autofocus lines into the pad, which was then topped with a 1 um thick carbon pad. A coarse trench was milled using a 30 nA FIB beam, stopping just before the pad’s front edge, and the trench face was subsequently fine polished with a 3 nA beam. SEM images were acquired at 1.5 kV accelerating voltage and a 1.1 nA beam current, with a pixel sampling of 4 nm, a slice thickness of 8 nm, and the in-lens EsB detector grid voltage was set at 820 V. The resulting image stacks were registered, inverted, and binned to generate volumetric reconstructions at 8 nm cubic voxel size.

To more easily and rapidly segment the rods in the FIB-SEM image volume, a model was trained in our napari plugin empanada^64^ using hundreds of 2D image patches containing the rods and the “train model” module (https://empanada.readthedocs.io/). This model was finetuned on additional images patches until it was reasonably able to segment rods in all three dimensions. Segmentation cleanup after running the model on the entire 3D volume was also done in napari. Additional segmentation of rod precursor structures was performed in 3DSlicer (Ver. 5.8.1;^68^) using the paint, erase, and level tracing segmentation tools. Movies of the FIB-SEM volume and rod segmentations were generated in Dragonfly (ORS; Ver. 2024.1 Build 1601). After all rod segmentations were finalised, successive dilate and fill-hole functions were applied to allow them to be modelled as solid cylinders. Rod lengths and diameters were then determined from the first and second principal components, respectively, of each of the cylindrical models. Rod measurements were then plotted using GraphPad Prism (Ver. 10.5.0 for Mac).

## Acknowledgements

We thank the Eickholt lab for support with equipment, the Liu lab for mouse tumour excision and fixation and the Edmondson lab for WT mouse lung excision and fixation. We thank Roland Wedlich-Söldner and Rebecca Eccles for critically reading the manuscript. We thank MHL for support with pathology and tissue preparation, particularly Jennifer Matta, Tamara Morgan, Thomas Sova and Sara Patamawenu for rod modelling and segmentation. We acknowledge the Advanced Medical Bioimaging Core Facility (AMBIO) of the Charité for support in acquisition and analysis of the imaging data. We thank the Advanced Light Microscopy & Image Analysis (ALM) facility of the Max-Delbrück-Center, the electron microscopy facility of the Max Planck Institute for Cell Biology and Genetics (MPI-CBG) and the technology platform of the Center for Molecular and Cellular Bioengineering (CMCB) for technical services. O.R. received funding by the Helmholtz Association, A.M. by the German Center for Diabetes Research and the German Ministry for Education and Research (BMBF).

## Author contributions

Conceptualisation, experimental design and supervision: O.R. and P.M.M. Conceptualisation of AT2 cell experiments: K.N. Investigations: E.B. (rod formation and dissolution), B.P., P.T. (temperature dependency, colocalisation studies), R.W.W. (IF experiments, BODIPY, cell lines), H.N., P.M.M. (temperature dependency, IF experiments, ATP depletion, OSER), S.A., F.H. (RHD protein experiments), S.K., P.M.M. (COS7 cell EM and CLEM), A.M. (EM tomography), K.R.L. (GP measurements), M.R.M. (AT2 cell EM), P.M.M. (OA and PA supplementation) O.R. (lipid metabolism inhibitors, cholesterol perturbation, RHD and ER protein localization, superresolution imaging, FRAP and LD experiments). Data analysis and visualisations: P.M.M. (most experiments). Manuscript writing: O.R., with critical editing from P.M.M., I.L., K.R.L., H.E. and K.N.

---

**Extended Data Fig.1.**
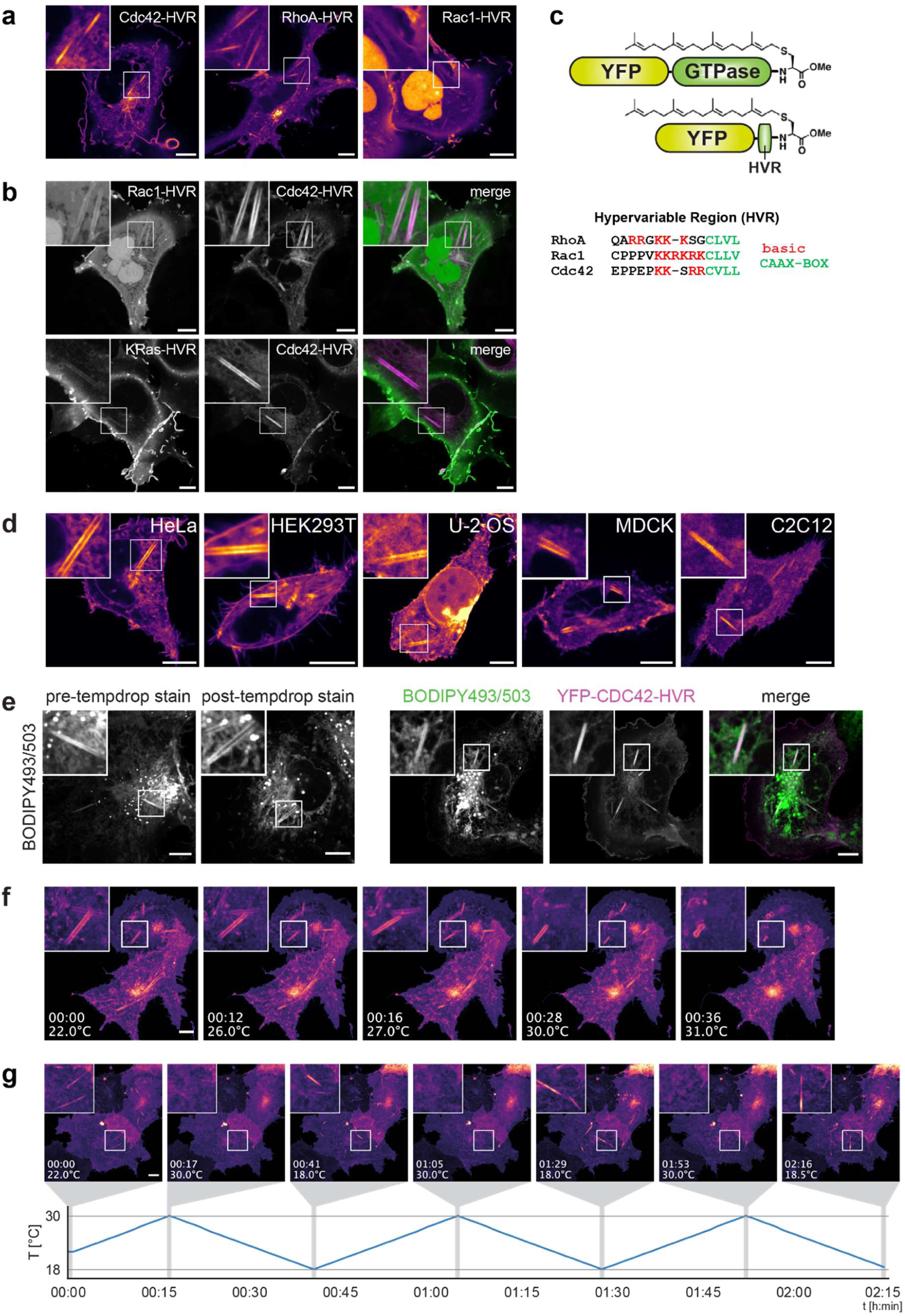
Rod markers, cell-line distribution, membranous nature and reversibility, related to Fig.1. **a,** Confocal images of live COS-7 cells incubated at 21°C for 40 min expressing the C-terminal hypervariable regions (HVR) of CDC42 (mCitrine-CDC42-HVR), RHOA (mCherry-RHOA-HVR) or RAC1 (mCherry-RAC1-HVR). All constructs carry geranylgeranyl lipid anchors. **b,** Co-expression of mCitrine-CDC42-HVR with mCherry-RAC1-HVR (top panel), or mCherry-CDC42-HVR with YFP-KRAS-HVR (bottom panel). YFP-KRAS-HVR carries a farnesyl lipid anchor. Insets highlight rod structures and differential enrichment of fluorescent reporters. **c,** Schematic comparison of full-length vs. HVR-only Rho GTPase constructs, indicating the geranylgeranyl lipid anchor and the corresponding HVR amino acid sequences. **d,** Live-cell confocal images of mCitrine-CDC42-HVR expressed in HeLa, HEK293T, U-2 OS, MDCK and C2C12 cells at 21°C, demonstrating consistent rod formation across cell types. **e,** Left: Confocal images of untransfected COS-7 cells labelled with the lipophilic dye BODIPY493/503 and incubated at 21°C for 40 min, demonstrating the membranous nature of rod structures. Staining was performed either before (pre-temp. drop) or after (post-temp. drop) cooling. Right: COS-7 cells expressing mCitrine-CDC42-HVR cooled to 21°C and subsequently labelled with BODIPY493/503, showing colocalization of both reporters. **f,** Time-lapse confocal images of COS-7 cells expressing mCitrine-CDC42-HVR. Rods present at 22°C (t=0) progressively shrink and release large vesicular structures during gradual rewarming to 31°C at 0.25°C/min. **g,** Reversible rod formation in COS-7 cells expressing mCitrine-CDC42-HVR during repeated thermal cycling between 21°C and 37°C, captured by time-lapse confocal imaging (corresponding to Supplementary Video4). Scale bars: 10 **-**µm. Time stamps in (**f**) and (**g**) are indicated in hh:mm.

**Extended Data Fig.2.**
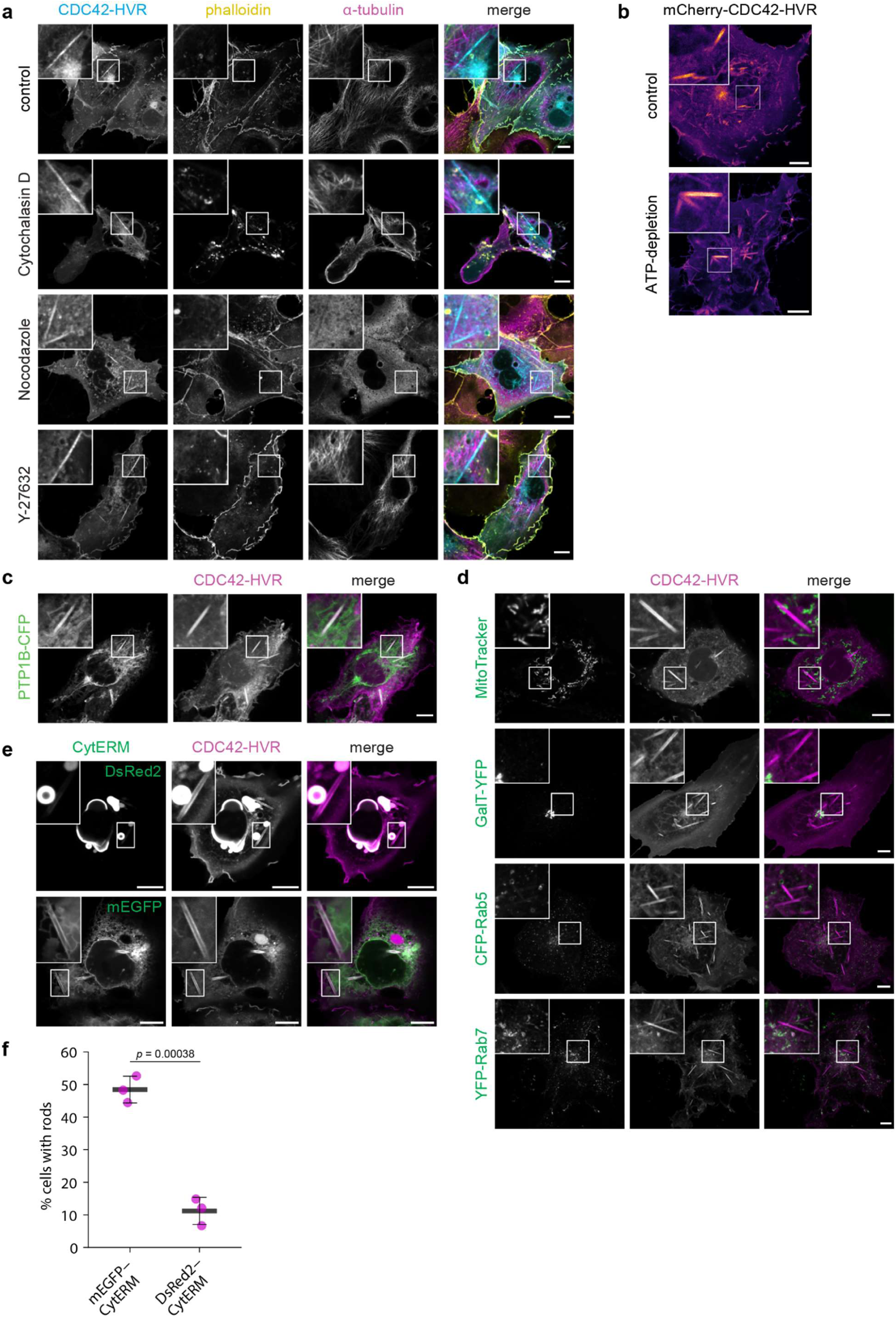
ATP- and scaffold-independent rod formation at the endoplasmic reticulum, related to Fig.1. **a,** Rod formation does not require structural support by the cytoskeleton and is unaffected by inhibition of cell contractility. COS-7 cells expressing mCitrine-CDC42-HVR were treated with Cytochalasin D (10 µM) or nocodazole (15 µM) for 30 min at 37°C, or the ROCK inhibitor Y-27632 (25 µM) for 10 min at 37°C. Cells were then equilibrated at 21°C for 40 min, fixed, stained for microtubules (anti-α-tubulin) and actin (phalloidin), and imaged by confocal microscopy. **b,** Rod formation does not require ATP. Live cell confocal imaging of COS-7 cells expressing mCherry-CDC42-HVR treated with or without ATP-depleting agents (0.05% sodium azide and 20 mM 2-deoxy-D-glucose) for 15 min at 37°C, followed by incubation at 21°C for 40 min prior to imaging. **c,** Live-cell confocal images of COS-7 cells coexpressing mCherry-CDC42-HVR and the ER marker PTP1B-CFP, an ER-localised peripheral membrane protein, after equilibration at 21 °C for 40 min. **d,** Confocal live images of COS-7 cells expressing mCitrine-CDC42-HVR labelled with MitoTrackerRed (top panel), or coexpressing mCherry-CDC42-HVR with the Golgi marker GalT-YFP (second panel), or coexpressing the vesicular compartment markers CFP-Rab5 with mCitrine-CDC42-HVR (third panel) or YFP-Rab7 with mCherry-CDC42-HVR (bottom panel). Cells were equilibrated at 21°C for 40 min prior to imaging. **e,** Ordered smooth ER (OSER) formation induced by a dimerising fluorescent protein fused to the cytoplasmic domain of a tail-anchored ER membrane protein (DsRed2-CytERM, top panel), whereas the monomeric version (mEGFP-CytERM) does not induce OSER. Rods still form in OSER-containing cells, albeit at reduced frequency, and exhibit a clearly distinguishable morphology. **f,** Quantification of rod formation in cells expressing mEGFP-CytERM versus DsRed2-CytERM shows reduced rod formation in the presence of OSER, likely reflecting limited availability of regular ER membrane material. Black crossbars indicate the mean of three independent experiments (magenta dots). Error bars: SD. Scale bars: 10 µm.

**Extended Data Fig.3.**
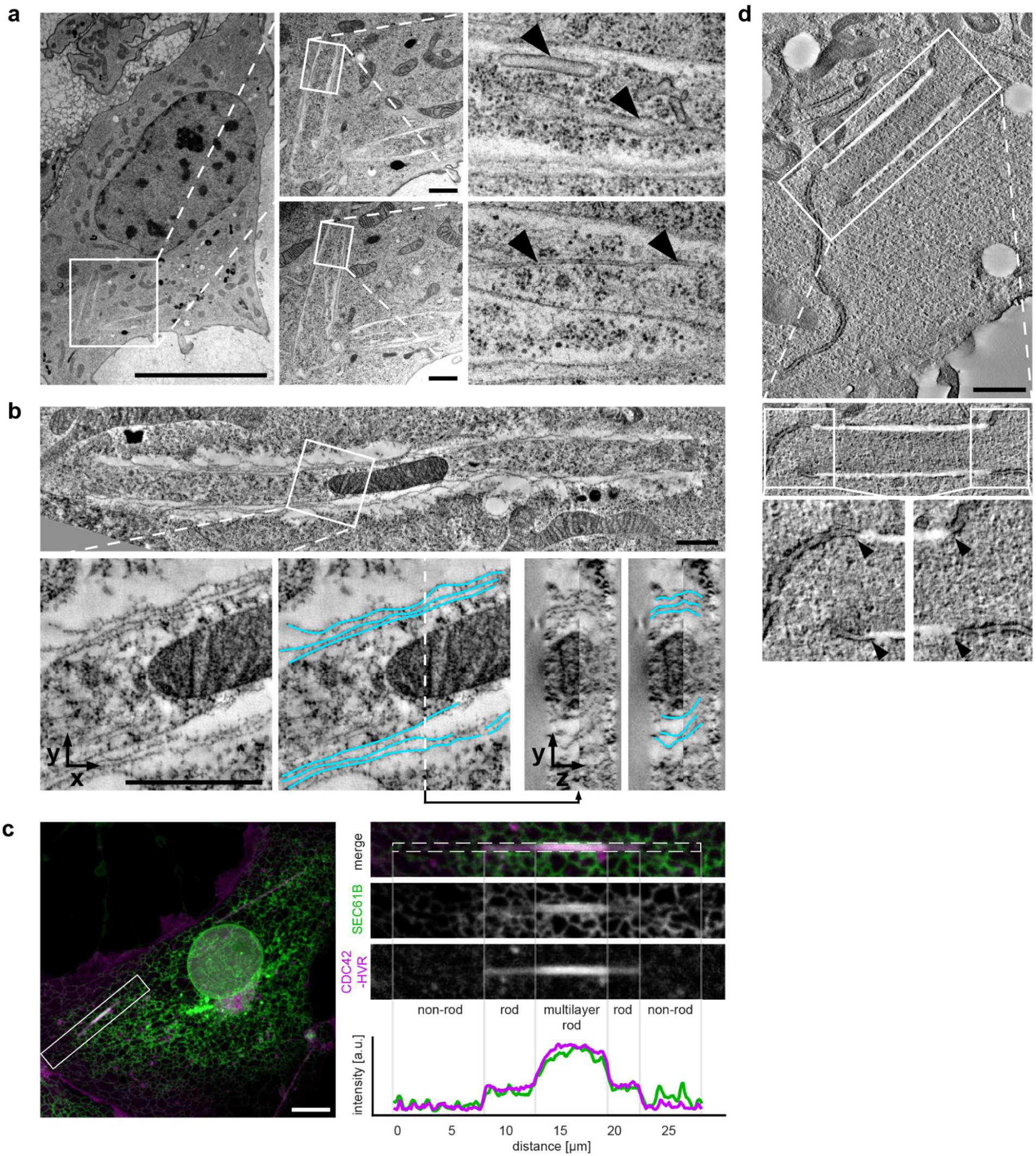
Ultrastructural analysis of rods, related to Fig.2. **a,** Left: Electron micrographs of high-pressure-frozen COS-7 cells confirming the rod morphology observed after chemical fixation in Fig.2. The boxed region is shown as two adjacent serial sections in the middle panels. Higher-magnification images of the boxed areas from these sections are shown on the right. Arrowheads in the upper right panel indicate ER membranes enclosed within the rod, arrowheads in the lower right panel mark microtubules within the same rod. Scale bars: 10 µm (left), 1 µm (centre). **b,** Top: Serial-section electron tomogram of high-pressure-frozen COS-7 cell showing a mitochondrion enclosed within a rod. Bottom left: Magnified view of the boxed region. Middle: Segmentation of rod membrane (blue), revealing its multilamellar organisation. The dashed lines marks the position of the orthogonal cross-section of the rod shown in right panels, further confirming the multilamellar architecture. Scale bars: 1 µm. **c,** Discrete, stepwise decline in fluorescence intensity occasionally observed towards rod termini in confocal images. Live-cell confocal images of COS-7 cell shown in Fig.1g and Supplementary Video5 coexpressing SEC61B-mEm (green) and mCherry-CDC42-HVR (magenta) after equilibration at 21°C for 40 min. The 30-min timepoint is shown. The boxed region is magnified on the right. Integrated fluorescence intensity within the dashed region is plotted below. Scale bar: 10 µm. Cells shown in the TEM panels were equilibrated at 21°C for 40 min prior to fixation or high pressure freezing. **d,** Electron tomogram of high-pressure-frozen COS-7 cell showing continuity between the rod and ER membrane at rod ends (black arrowheads). Scale bar: 1 µm

**Extended Data Fig.4.**
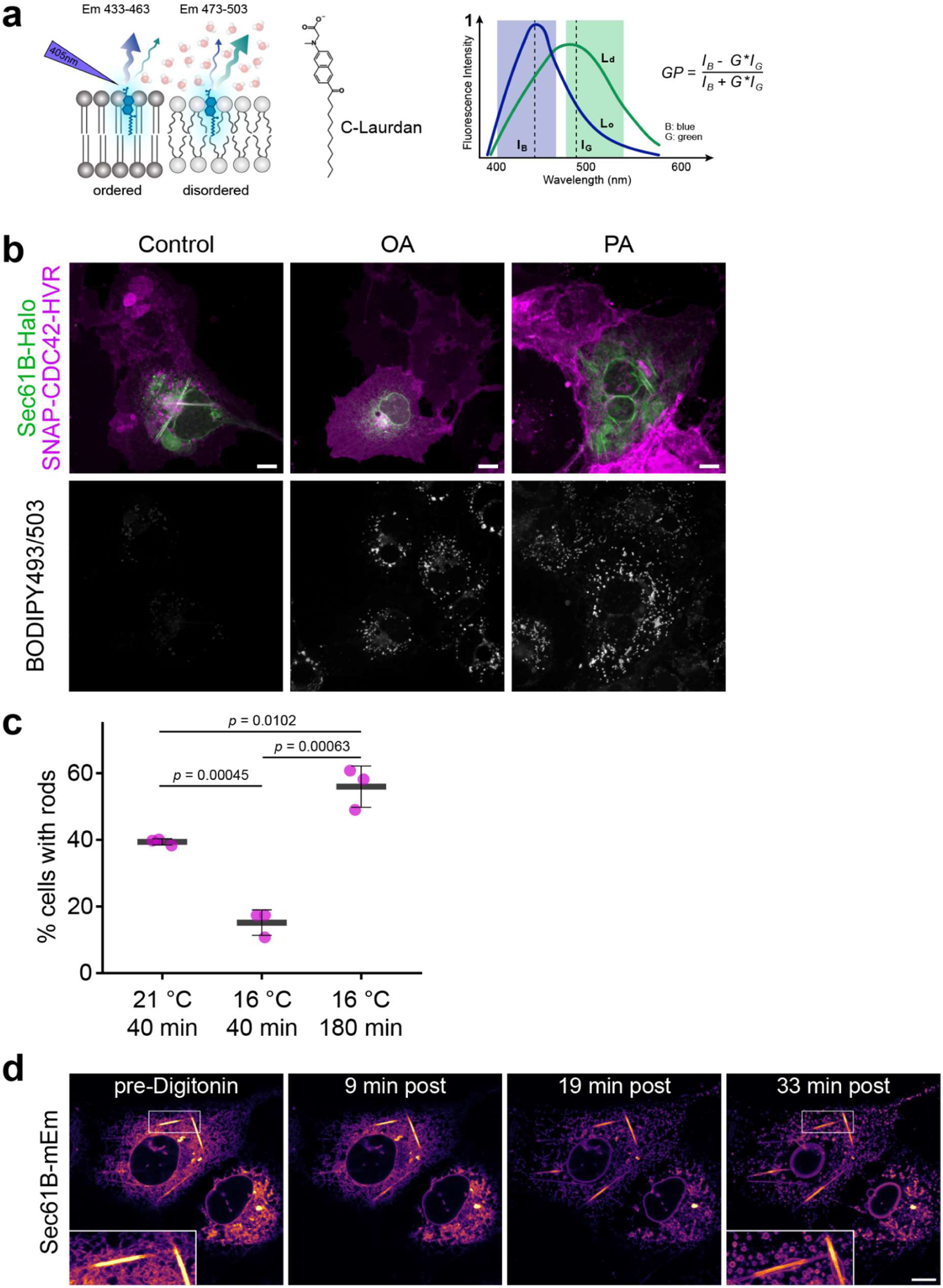
Fatty-acid-induced lipid droplet formation, diffusion-limited phase separation, and rod detergent resistance, related to Fig.3. **a,** Schematic illustration of General Polarisation (GP) measurements using the solvatochromic probe C-Laurdan. In the GP equation, IB: represents the intensity in the blue channel, I_G_ the intensity in the green channel and G the calibration factor. **b,** Confocal images of COS-7 cells expressing JF646-labelled SEC61B-Halo and JF-549-labelled SNAP-CDC42-HVR after 48h treatment with oleic acid (OA; 50 µM) or palmitic acid (PA; 50 µM). Cells were stained with BODIPY493/503 for 1 h and subsequently cooled to 21°C for 40 min prior to imaging. Scale bars: 10 µm. **c,** Percentage of rod-positive CDC42-HVR-expressing cells after cooling to the indicated temperatures for the indicated durations. Cooling below 21°C has opposing effects: it increases membrane viscosity and promotes rod formation. At fixed incubation times, higher viscosity restricts lipid diffusion and domain coalescence, reducing rod formation. With longer incubation, the stronger low-temperature phase-separation drive dominates, enhancing rod formation. Black crossbars indicate the mean of three independent experiments (coloured dots). SD shown. **d,** Confocal time-lapse images of COS-7 cells expressing SEC61B-mEm maintained at 21°C and treated with digitonin (10 µM) for the indicated times. Digitonin disrupts cellular endomembranes but does not cause rod collapse. Scale bars: 10 µm.

**Extended Data Fig.5.**
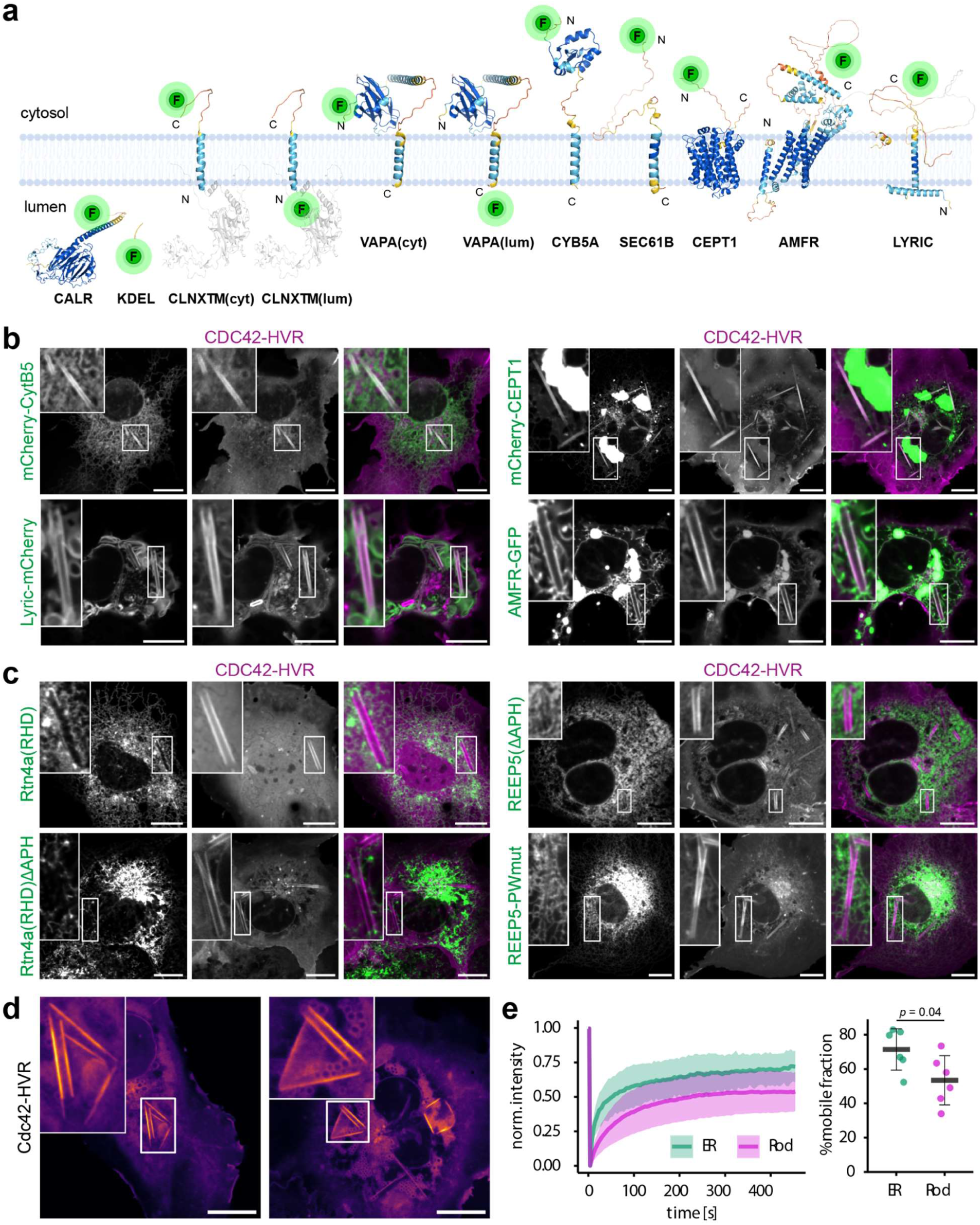
Protein exclusion and diffusion within rods, related to Fig.4. **a,** Membrane topology and fluorescent tag-orientation of the ER-resident proteins used in this study. Structures are based on AlphaFold predictions. Parts of the disordered regions of LYRIC (193-398 and 462-end) and of Calnexin (CLNX, aa 1-15 and 523-end) were omitted for clarity of display. The truncated luminal N-terminus of Calnexin absent from the N-terminally and C-terminally tagged constructs are greyed out. **b,c,** Representative confocal micrographs of COS-7 cells expressing the indicated plasmids after cooling to 21°C for 40 min. Localisation on rods versus exclusion from rods: mCherry-CytB5 (35/0), Lyric-mCherry (17/0), mCherry-CEPT1 (21/0), AMFR(gp78)-GFP (20/5), GFP-RTN4A-RHD (RHD only) (0/30), GFP-RTN4-RHD-ΔAPH (0/46), mScarlet-REEP5-ΔAPH (0/19), REEP5-PWmut-mEm (P63L/W84R) (0/15) **d,** Examples of rod-structures connected to giant membrane sheets with sharp linear edges, potentially representing stalled intermediates during conversion into fully wrapped multilamellar rods. **e,** COS-7 cells expressing JF646-labelled SEC61B-Halo and JF549-labelled SNAP-CDC42HVR were cooled to 21°C for 40 min and subjected to fluorescence recovery after photobleaching (FRAP). Left: normalised fluorescence intensity in photobleached regions over time (n = 6 cells per region, error margins indicate SD). Right: mobile fraction calculated from the final 5 time points (430-450s post-bleach), error bars indicate SD. Scale bars: 10 µm.

**Extended Data Fig.6.**
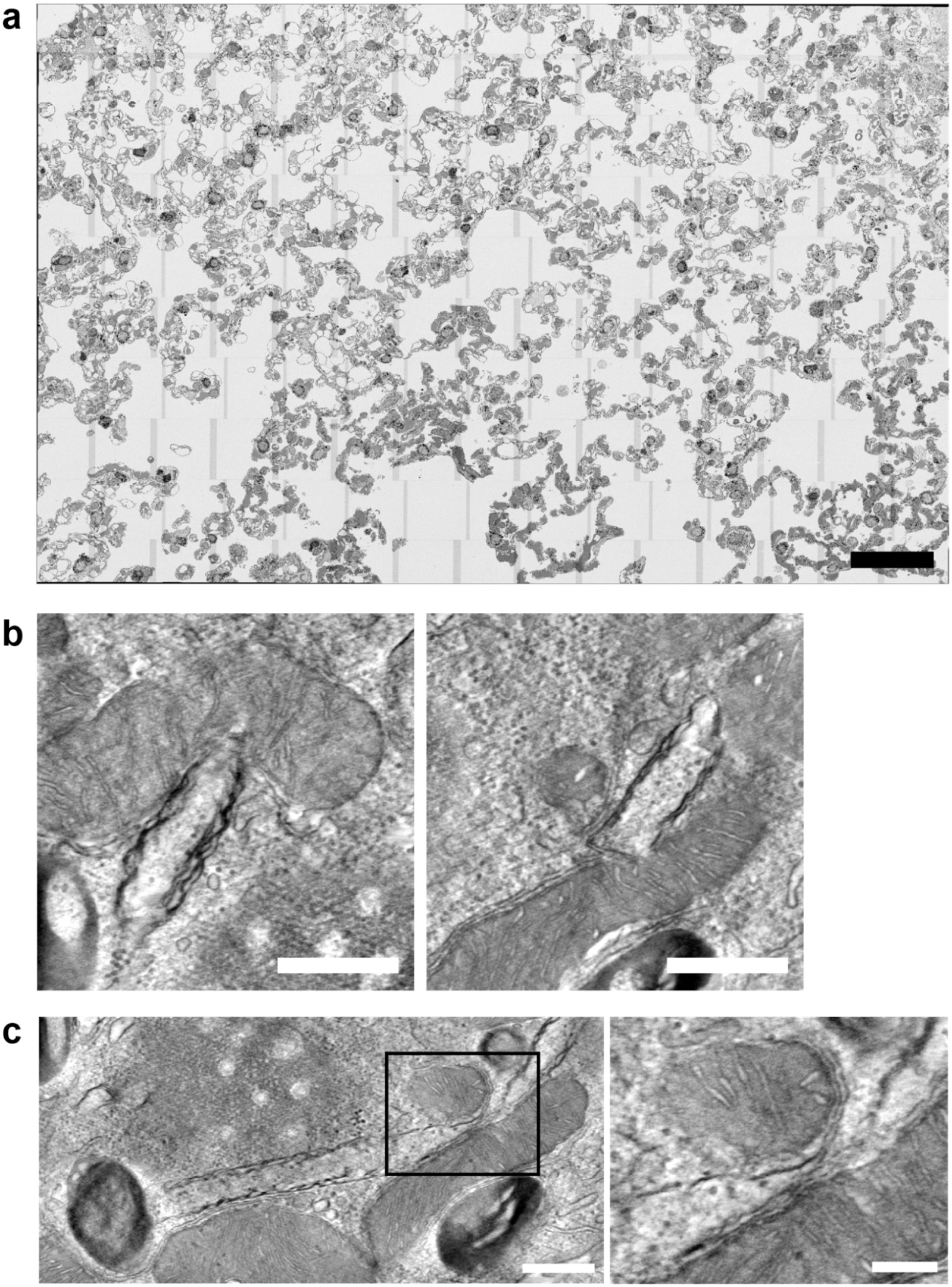
Electron microscopy of mouse lung AT-2 cells, related to Fig. 5. **a,** Overview of the full imaging area of the *Kras^LA^*^2^ lung tumour sample fixed at 21°C, corresponding to Fig.5a. **b,** TEM tomographic slices showing rods deforming mitochondria. **c,** Evidence of membrane continuity between rods and ER. The boxed region is shown at higher magnification on the right. Scale bars 50 nm (**a**), 500 nm (**b,c** left), 200 nm (**c,** right).

**Supplementary Video 1** Confocal time-lapse movie of COS-7 cells expressing mCitrine-CDC42-HVR cooled from 37°C to 20°C at 0.5°C/min. Scale bar: 10 µm. Time stamps indicate hh:mm (corresponding to Fig.1e).

**Supplementary Video 2** Confocal time-lapse movie of COS-7 cells expressing mCitrine-CDC42-HVR which formed rods at 22°C before warming to 31°C at 0.25°C/min. Scale bar: 10 µm. Time stamps indicate hh:mm (corresponding to Fig.1f).

**Supplementary Video 3** Confocal time-lapse movie of COS-7 cells expressing mCitrine-CDC42-HVR which formed rods at 22°C before warming to 31°C at 0.25°C/min. Scale bar: 10 µm. Time stamps indicate hh:mm (corresponding to Extended Data Fig.1f).

**Supplementary Video 4** Confocal time-lapse movie of COS-7 cells expressing mCitrine-CDC42-HVR subjected to repetitive cooling and heating cycles between 18°C and 30°C. Scale bar: 10 µm. Time stamps indicate hh:mm (corresponding to Extended Data Fig.1g).

**Supplementary Video 5** Confocal time-lapse movie of COS-7 cells expressing SEC61B-mEmerald (green) and mCherry-CDC42-HVR (magenta), instantaneously cooled to 21 °C, revealing colocalization of both proteins during rod formation and elongation. Time stamp indicates time after cooling to 21°C. Magnified views reveal intensity steps along the rod (corresponding to Extended Data Fig.3c). Note the stalled sheet-rod intermediate (corresponding to Extended Data Fig.5d) in the lower right, followed by a sheet-to-rod transition (corresponding to Fig.4e and Supplementary Video11). Scale bar: 10 µm.

**Supplementary Video 6** Confocal time-lapse movie of COS-7 cells expressing SEC61B-mEm and mCherry-CDC42-HVR, cooled to 21°C, showing small peripheral rods connected to the ER network.

**Supplementary Video 7** Deconvolved 3D super-resolution image of a COS7 cell expressing JF646-labelled SEC61B-Halo, cooled to 21°C, highlighting rod morphology and connection to surrounding ER (corresponding to Fig.1i).

**Supplementary Video 8** Serial-section electron tomography of a rod cross-section and 3D segmentation showing the rod membrane (blue) and its continuity with the ER network (yellow) (corresponding to Fig.2e,f).

**Supplementary Video 9** Confocal time-lapse movie of COS-7 cells expressing SEC61B-mEmerald cooled to 21°C, showing persistence of rods during apoptotic cell shrinkage and morphological collapse.

**Supplementary Video 10** Confocal time-lapse movie of a COS-7 cell expressing GFP-RTN4A (green) and mCherry-CDC42-HVR (magenta) maintained at 21°C, showing accumulation of RHD-containing protein at end-to-end contacts between ER tubules and rods.

**Supplementary Video 11** Confocal time-lapse movie of a COS-7 cell expressing JF646-labelled SEC61B-Halo (green) and JF-549-labelled SNAP-CDC42-HVR (magenta) maintained at 21°C, showing sheet-to-rod transition.

**Supplementary Video 12** Deconvolved 3D super-resolution projection of a COS7 cell expressing JF549-labelled SNAP-CDC42HVR maintained at 21°C, highlighting a stalled sheet-to-rod intermediate (corresponding to Extended Data Fig.5d).

**Supplementary Video 13** Three-dimensional FIB-SEM volume of an AT-2 cell (corresponding to Fig.5b) from a *Kras^LA^*^2^ lung tumour fixed at room temperature. Individual rods were segmented in Empanada (napari) and rendered in distinct colours.

**Supplementary Video 14** FIB-SEM volume corresponding to Fig.5j showing a putative sheet-to-rod intermediate. The structure appears as a hollow rod enclosing a mitochondrion at the lower right and extends toward the nucleus, where it adopts a sheet-like morphology. Additional rods are present within the volume, several of which also enclose mitochondria. Scale bar: 1 µm.

**Supplementary Video 15** Tomographic reconstruction corresponding to Extended Data Fig.6c of an AT-2 cell from a *Kras^LA^*^2^ lung tumour fixed at room temperature, showing continuity between rods and morphologically regular endoplasmic reticulum. Scale bar: 500 nm.

